# Integrating multiple data sources to fit matrix population models for interacting species

**DOI:** 10.1101/594986

**Authors:** Frédéric Barraquand, Olivier Gimenez

## Abstract

Inferring interactions between populations of different species is a challenging statistical endeavour, which requires a large amount of data. There is therefore some incentive to combine all available sources of data into a single analysis to do so. In demography and single-population studies, Integrated Population Models combine population counts, capture-recapture and reproduction data to fit matrix population models. Here, we extend this approach to the community level in a stage-structured predator-prey context. We develop Integrated Community Models (ICMs), implemented in a Bayesian framework, to fit multispecies nonlinear matrix models to multiple data sources. We assessed the value of the different sources of data using simulations of ICMs under different scenarios contrasting data availability. We found that combining all data types (capture-recapture, counts, and reproduction) allows the estimation of both demographic and interaction parameters, unlike count-only data which typically generate high bias and low precision in interaction parameter estimates for short time series. Moreover, reproduction surveys informed the estimation of interactions particularly well when compared to capture-recapture programs, and have the advantage of being less costly. Overall, ICMs offer an accurate representation of stage structure in community dynamics, and foster the development of efficient observational study designs to monitor communities in the field.

## 1 Introduction

Although matrix population models (MPMs) are well developed for single-species, they have received limited attention when it comes to modelling the intricate dynamics of interacting species. Nonlinear matrix population models (Cushing, 1988; Neubert & Caswell, 2000), which admit a standard MPM formulation at low densities, are density-dependent models with nonlinearities possibly incorporating interactions between classes (e.g., stages or species) as functions of the abundances of these classes (Hassell & Comins, 1976; Travis *et al*., 1980; Cushing, 1988; Dennis *et al*., 1995). However, outside the celebrated study of the floor beetle dynamics (Dennis *et al*., 1995), nonlinear MPMs have been little used to estimate interactions between species or reproduce their joint dynamics.

Nonlinear MPMs could, however, contribute greatly to ecology. Strongly fluctuating dynamics are currently believed to be as often driven by stage structure than by predator-prey interactions, with possible interactions between both mechanisms (Murdoch *et al*., 2002; de Roos & Persson, 2013). Fitting nonlinear MPMs to data would allow ecologists to better understand the relative share and potential interactions of those mechanisms in generating population fluctuations (Barraquand *et al*., 2017). Another leading ecological question is how do many species coexist in spite of seemingly similar ecological requirements: nonlinear MPMs allow one to distinguish competition between several life-stages (Fujiwara *et al*., 2011), hence paving the way for a better understanding of coexistence than unstructured models (e.g., there could be much stronger intraspecific density-dependence at the seedling stage and yet similar intra-vs. inter-specific competition at the adult stage, Chu & Adler, 2015). Stage structure may also affect the strength of trophic cascades and has been proposed to be one of the main developments needed in community-level models (Miller & Rudolf, 2011). Given all this potential, there is reason to reflect on the relative paucity of stage-structured (or age-structured) community-level models fitted to data (but see Adler & HilleRisLambers 2008; Chu & Adler 2015 in plants). There is a literature on stage-structured consumer-resource studies in continuous-time (de Roos & Persson, 2013), yet it is largely theoretical.

The scarcity of empirically-parameterized, multi-species and nonlinear MPMs may partly be due to their increased dimensionality. Indeed, a system with interactions between *S* species and *L* stages per species requires estimation of (*S × L*)^2^ interaction parameters; this may be why unstructured statistical models for interaction between species have so far been preferred (Ives *et al*., 2003), at least when a single type of data is used (e.g., time series of counts, Dennis *et al*. 1995).

Although nonlinear MPMs have many parameters, because of their internal age-structure, they also have advantages over unstructured discrete-time models currently fitted to data (i.e., discrete-time Lokta-Volterra or log-linear autoregressive modelling; Ives *et al*. 2008; Mutshinda *et al*. 2009; Hampton *et al*. 2013). Indeed, the survival rates expressed in MPMs are well estimated by capture-recapture techniques (Caswell, 2001; Lebreton *et al*., 2009). This opens new avenues to fit nonlinear matrix models for multiple species, by considering other types of data than just counts of species, such as data on survival and development rates, as well as reproduction. One approach to incorporate such demographic datasets, used in plant community dynamics, is to fit separate models for reproduction and survival components of the demography (Adler & HilleRisLambers, 2008; Chu & Adler, 2015), and then simulate the community-level model thus created, to evaluate its prediction of the counts and spatial structure. While this approach is sound, it does not take full advantage of opportunities to combine vital rate data with counts, which might be problematic for small datasets.

Capture-recapture and reproduction data can be combined advantageously with counts within the Integrated Population Modelling framework (Besbeas *et al*., 2002), which uses MPMs as its core for integrating over several datasets through products of likelihoods. This framework has recently been extended to density-dependent MPMs (Abadi *et al*., 2012). To the best of our knowledge, there has been only one comprehensive attempt to move integrated modelling from the population to the community level while including species interactions (i.e., density-dependent links from species *j* density to species *i* vital rates), that of Péron & Koons (2012). Other multi-species IPMs have been recently published, but they do not model interactions (e.g., Lahoz-Monfort *et al*., 2017) or include interactions, but model the demography of one species only (e.g., Saunders *et al*., 2018).

Although Péron & Koons (2012) did provide a proof of concept for community-level integrated modelling, their focus was on estimating parameters in a real two-species duck community in order to understand species coexistence, not performing a general assessment of the relevance of integrated community models (ICMs). The value of the different sources of data and combining them into ICMs has therefore not yet been evaluated, which is what we attempt here. To do this, we start by developing a theoretical model, which we then simulate under different scenarios, to find out which combination of data sources allows the model to be identifiable in practice. Our model also differs from Péron & Koons (2012) in that it models predator-prey interactions rather than competition or parasitism, thus occupying a new and complementary niche in the vast space of (under-developed) density-dependent MPMs. We used the literature on nonlinear matrix models (Cushing, 1988; Neubert & Caswell, 2000; Wikan, 2001; Fujiwara *et al*., 2011) as an inspiration for model development, and we therefore produced a model that is not only statistically-friendly, but also amenable to theoretical explorations (i.e., it is permanent *sensu* Kon *et al*. 2004; Benaïm & Schreiber 2009, for a large swath of parameter space).

In the following, we first formulate a nonlinear matrix model to represent predator-prey interactions between two species, inspired by earlier work on stage-structured density-dependent matrix models. Our model therefore connects to Neubert & Caswell (2000) (single-species density-dependent models) as well as Dennis *et al*. (1995), who modelled cannibalism within a single species. We then bring nonlinear matrix models into a statistical framework combining multiple data sources instead of only counts.

Our work has two main ramifications for population and community ecology. From a theoretical standpoint, we pave the way for a theory of predator-prey interactions based on demography rather than biomass flow. This takes into account not only direct effects of predation, but also indirect effects that have been neglected for a long time (Preisser *et al*., 2005). From a more empirical perspective, we weight the information brought by the different sources of data that can be collected in the field, which has major implications for study design: when initiating a community-level survey, should one prioritize capture-recapture, reproduction or count data?

## 2 Models

The two-stage predator-prey model studied here belongs to the class of models obeying the life-cycle graph shown in Fig. 1, whereby adult predators negatively affect juvenile prey survival and any increase in juvenile prey density positively impacts predator fecundity.

**Figure 1:**
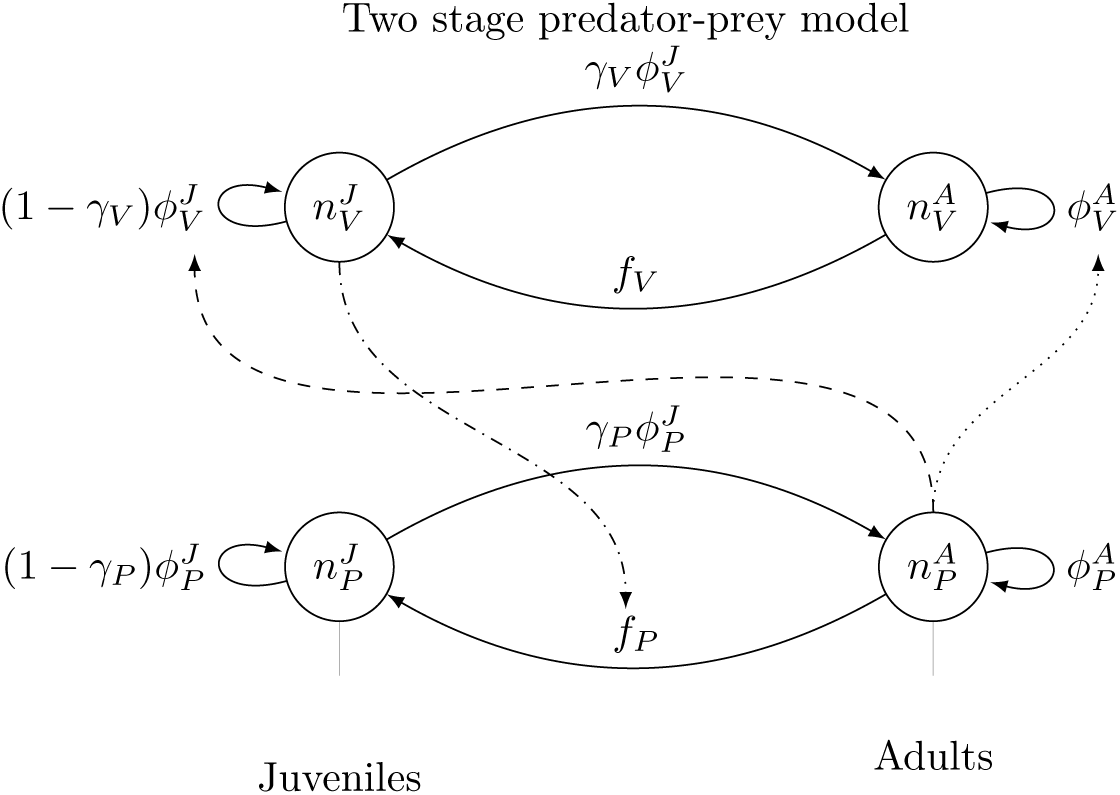
Life cycle graph with added interspecific interactions. Plain lines describe transitions within the prey or the predator population, while dashed lines describe which parameters (pointed by the arrowhead) are affected by which density (origin of the arrow). In our main model, juvenile prey affects the predator reproduction and predator density affects juvenile prey survival (logical for bird/mammal predators eating bird/mammal youngs). The dotted line represent a model variant where predation would also affect adult prey survival. In all the models fitted in this paper, γP = γV = 1.

Predators are denoted by the symbol *P* and prey by *V* (as in ‘victim’). We chose to model an interaction where the predation by adult consumers affects the juvenile stage of the prey, as this is common in the systems we have in mind (birds and mammals, for which count, reproduction and capture-recapture data are readily available). Our ‘target system’ could feature a mammal or bird top predator eating ground-breeding bird chicks (as often do canids or birds of prey, Valkama *et al*. 2005) or a carnivorous mammal eating preferentially juvenile ungulates (e.g., 3 carnivores out of 4 in Gervasi *et al*., 2012). This adult predator - juvenile prey interaction setting promotes rather mild fluctuations in numbers for the parameters that we considered (as opposed to say, predation on adults without a refuge, generating a full-blown limit predator-prey cycle, e.g. Turchin & Hanski 1997). This setting is all the more relevant to test our ICM framework that interactions between predators and prey will be harder to detect from time series alone in the absence of large oscillations. Time series techniques for mechanistic predator-prey models perform well for cyclic or chaotic dynamics (e.g., Turchin & Ellner, 2000), but it is not quite clear that model identification is always possible for milder dynamics from counts only.

It must be mentioned, however, that for other parameters than those considered here (we tuned them to slow life-histories of birds and mammals), density-dependent models can and do produce cycles, resonances, and chaos (Neubert & Caswell, 2000; Greenman & Benton, 2004). This is especially likely for very large maximal fecundities (see Discussion and Appendix A), which may impact the estimation process.

For simplicity, we set the maturation rates of juvenile prey (*γ*_*V*_) and predator (*γ*_*P*_) to 1, so that juveniles of the year mature at the end of one time period to adulthood. Note that if the maturation time is larger than a year, the use of a different time unit (e.g., 3 years) may allow one to use this model parameterization even though *γ*_*P*_ = *γ*_*V*_ = 1 (aggregating yearly reproduction into larger time intervals may be more or less appropriate depending on the speed of life histories). Other models with *γ <* 1 could be generalized to a stochastic ICM, but are somewhat more complex to simulate and fit (see Discussion and Appendix E).

In the following, we first describe the baseline nonlinear, stage- and species-structured matrix population model that forms the (deterministic) backbone of our work. Then, we introduce stochasticity, due to individual discreteness, and formulate the Integrated Community Model that will be fitted to the data.

### 2.1 Nonlinear matrix model with predator-prey and stage structure

The baseline predator-prey model that we consider can be expressed as

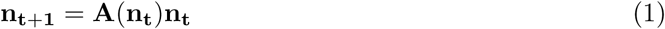

for the abundances of the two species and the two stages per species, with

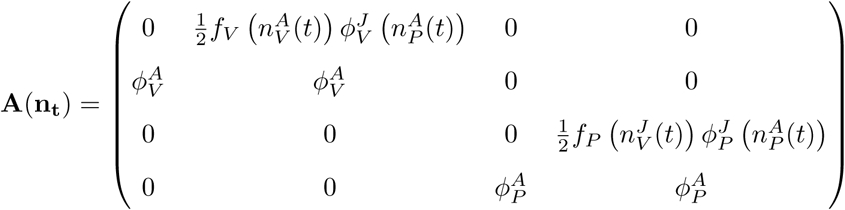

the population projection matrix and the population abundance vector

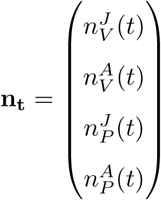

This model is a pre-breeding model, which assumes that population census is taken right before reproduction (Cooch *et al*., 2003). We assume that all adults, and only adults, are able to reproduce and can do so every year.

We use the following notations:

- 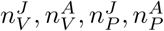 are the abundances of juvenile prey, adult prey, juvenile predators and adult predators, respectively.
- *f*_*V*_, *f*_*P*_ are the number of prey juveniles (respectively predator juveniles) produced by a prey adult (resp. predator), during one time unit.
- 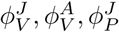 and 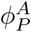 are the fractions of the juvenile prey, adult prey, juvenile predator and adult predator surviving between *t* and *t* + 1.

The model encapsulates the following ecological relationships:

- Prey reproduction (*f*_*V*_) decreases with the number of adult prey individuals
- Juvenile prey survival 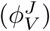 decreases with adult predator density
- Predator juvenile survival 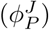 decreases with adult predator density
- Predator reproduction (*f*_*P*_) increases with juvenile prey numbers

These relationships are modelled with the following functions:

- 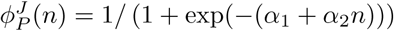
- 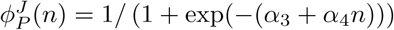
- *f*_*p*_(*n*) = exp(*α*_5_ + *α*_7_*n*)
- *fv (n)* = exp(*α*_7_ + *α*_*s*_*n*)

The exponential function for fecundity (log link function) is classical for IPM modelling (Schaub & Abadi, 2011) and maintains a much-needed connection to theoretical work on such density-dependent models (Neubert & Caswell, 2000). Although a sigmoid-shaped survival rate makes empirical sense, this choice was motivated by statistical estimation purposes. Identical choices of functional forms have been made by Péron & Koons (2012).

For the parameter values that we consider (Table 5), representing mammal or bird demographies, the deterministic model produces a stable fixed point (Appendix A). The intraspecific density-dependent feedbacks have been included so as to produce a relatively stable model with minimal complexity. By stable, we mean here that the model converges over time either towards a stable fixed point or a stable ‘manifold’ (some attractor being a *k*-point cycle, an invariant loop, or a strange attractor; see e.g., Caswell 2001, chapter 16), where trajectories are bounded when time tends to infinity. Only considering interspecific density-dependent links, by contrast, would lead instead to an unstable model, reminiscent of the behaviour of the Nicholson-Bailey model (also in discrete-time and without any intrinsic regulation between prey species; Kot 2001, chapter 11). In other words, without intra-specific density-dependence, the model trajectories would eventually ‘blow-up’ and head towards zero or infinity (Appendix A): simulating from the fitted model, for instance, would become meaningless and interaction estimates would be hard to interpret.

A stable fixed point in a multispecies model is usually a scenario where the low amount of fluctuations in the time series generates in turn little information in the count data alone. This is therefore a parameter scenario that is both realistic for the study system we have in mind (birds and mammals with slow dynamics) and relevant to incorporate capture-recapture and reproduction together with count data. Cases where the deterministic age-structured model itself produces a limit cycle, an invariant loop, or strange attractor (Neubert & Caswell, 2000) are left for further work.

That said, one should keep in mind that all forms of stochasticity, including demographic stochasticity, do interact with the deterministic skeleton of the model, which has been especially well shown in predator-prey models (Nisbet & Gurney, 1976; McKane & Newman, 2005). Therefore, a stable fixed point in the deterministic model does not guarantee that the dynamical behaviour of a stochastic model will consist only of small fluctuations around the fixed point. Although this seemed to be the case for the first parameter set that we considered (see below), the second showed more variability, with some oscillations when simulated for long times (Appendix A).

### 2.2 Stochastic Integrated Community Model

Here, we first highlight the population-level predator-prey model with added demographic stochasticity, which produces discrete-valued counts. We then add the statistical part of the capture-recapture and reproduction submodels that require some individual-based definitions, and are overlayed on top of the stochastic MPM. It is the general combination of the dynamic process model with the count model, the capture-recapture model, and the reproduction model that makes the Integrated Community Model (ICM). The ICM was both simulated and fitted by Monte Carlo Markov Chain (MCMC) methods in JAGS (Plummer *et al*., 2003). The computer code is provided at https://github.com/oliviergimenez/predator_prey_icm.

#### 2.2.1 Stochastic multispecies nonlinear model

Now we consider a discrete-valued state vector **n**_*t*_. At each time step, adult predators survive with probability 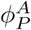, together with juveniles of the previous year (pre-breeding census, eq. 1), so that

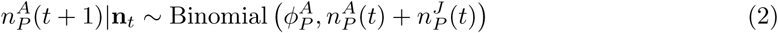

which is just a Binomial sampling of eq. 1. The expected mean number of juveniles produced by an adult at *t* + 1, *m*_*P*_ (*t* + 1), combines adult fecundity *f*_*P*_ and newborn survival through the year with probability 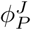 both of which are function of **n**_*t*_, so that 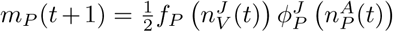 (see eq. 1). The number of juveniles is then updated as

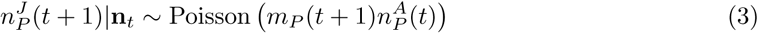

The equations are similar for prey counts

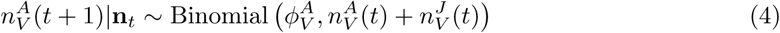

with 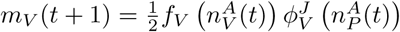 (see eq. 1). The number of juveniles at *t +* 1 is then

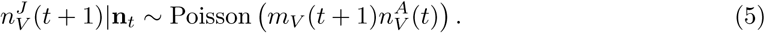

This ends the mechanistic (dynamic process model) part of the stochastic ICM.

#### 2.2.2 Statistical part of the ICM

The ICM combines data from counts, capture-recapture, and reproduction, which implies a statistical model for each.

The model for counts assumes that juveniles and adults are aggregated (not distinguishable). We overimpose an additional observation error (customary in population ecology), which leads to the equation:

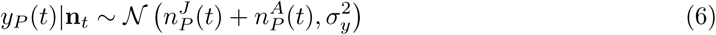

The equation for the prey is obtained by symmetry. In practice, this observation error is set to a very small value (Table 5) because of issues with unknown count observation error (see Discussion).

The capture-recapture model is a standard Cormack-Jolly-Seber model with two age classes (see e.g., Lebreton *et al*., 1992; Kéry & Schaub, 2011, for a detailed exposition). The model for fecundity (also called productivity in bird studies) is a Poisson regression: the total number of newborns counted in year *t* follows a Poisson distribution with rate *R × f* where *R* is the number of surveyed broods and *f* the fecundity. This assumes that each mother produces a Poisson distributed number of offspring, so that any sum of reproductive output over a number of mothers is also Poisson distributed.

Each model was fitted using two MCMC chains, 10000 iterations for burn-in and 20000 additional time steps to ensure convergence in the baseline scenario.

All priors were taken as vague (flat) or weakly informative (e.g., standard Gaussian), in order to provide an estimation setting not too dissimilar to frequentist approaches (our goal here was to develop ICMs, not to favor any inferential framework for hierarchical models), and to be able to assess easily a first aspect of identifiability: a relatively flat prior allows one to check graphically if there is overlap between prior and posterior, which is a main proxy for identifiability in a Bayesian context (Gimenez *et al*., 2009). More specifically, all a priori probabilities of capture and constant probabilities were taken as Unif(0, 1). We used a priori *α*_*i*_ *∼ N* (0, 1). That being said, in extensions of this model for which the combination of data sources still led to substantial bias in estimates, we also considered more informative priors (see Discussion).

## 3 Simulation study

In order to show the potential benefits of combining data into Integrated Community Models, we fitted ICMs in four contrasted scenarios of data availability. We constructed those to mimic what is observed in the field, where count data is usually more available (Schaub & Abadi, 2011) even though less informative of vital rates directly (Zipkin & Saunders, 2018). There is also often more demographic information on longer-lived, large predators than on their prey, an asymmetry in data availability which we incorporated here, although we did not try to reproduce the observational data design of any species pair in particular.

Our data scenarios are as follows:

1. All data available for both predator and prey (Table 1)
2. Only counts available for both species (Table 2)
3. Reproduction missing for prey (Table 3)
4. Capture-recapture missing for prey (Table 4)

The scenario 1 helps to verify that the statistical model is working properly when all the information is available.

**Table 1:**
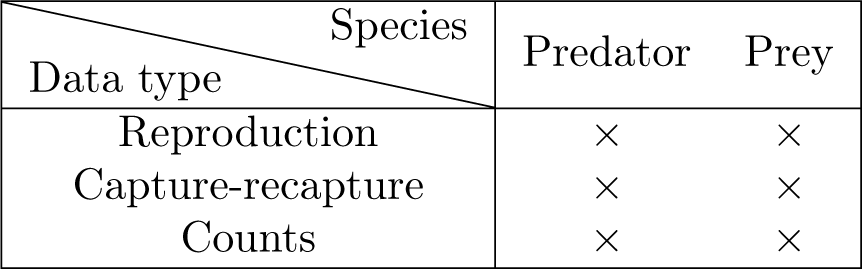
Scenario 1 - All data types available for both species

**Table 2:** Scenario 2 - Only counts, for both species

| Data type \ Species | Predator | Prey |
| --- | --- | --- |
| Reproduction |  |  |
| Capture-recapture |  |  |
| Counts | × | × |

Scenario 2 is then helpful to pinpoint identifiability issue whenever only counts are available.

**Table 3:** Scenario 3 - All on predator, counts for both, and capture-recapture for prey

| Data type \ Species | Predator | Prey |
| --- | --- | --- |
| Reproduction | × |  |
| Capture-recapture | × | × |
| Counts | × | × |

In scenario 3, predators are assumed to be easier to monitor, both species undergo capture-recapture, whilst prey reproduction is unknown.

**Table 4:**
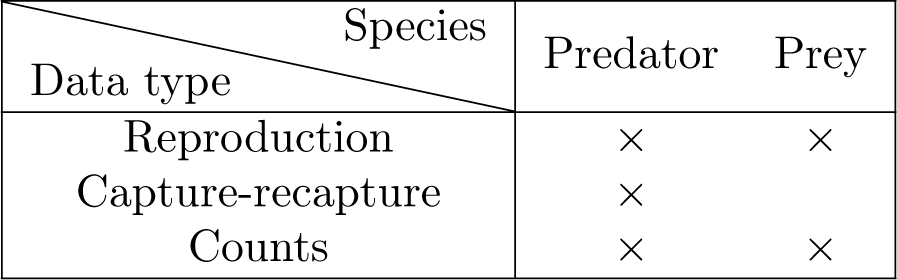
Scenario 4 - All on predator, counts for both, and reproduction data for prey

In scenario 4, predators are assumed to be easier to monitor, whilst prey capture-recapture data are not possible to obtain. Comparison of the last two scenarios will be useful to assess the value of a costly capture-recapture program for prey monitoring. The four scenarios are crossed with two population monitoring durations, *T* = 10 years and *T* = 30 years (assuming the time unit is a year). We assumed that 100 juvenile individuals are marked each year for *T* = 10, and 20 juvenile individuals are marked each year for *T* = 30, for both species. Therefore, *T* = 30 is not merely ‘more data’ but a different observational design. The capture-recapture study design implicitly assumes that it is easier to mark the predator, because they are less numerous than prey and the numbers of juveniles marked therefore represent a higher fraction of the predator population. The annual number of surveyed broods in set to 20 for the predator and 50 for the prey, hence, for both species, fecundity is sampled only for a fraction of the population. Table 5 describes all other models parameters that have been used for simulation.

For scenarios 2 and 4, the fitted models are without capture-recapture (in scenario 4, only for the prey): we kept a probability of capture in the code but it has nearly equal priors and posteriors and cannot affect other estimates (it only enters the capture-recapture submodel, which is absent).

**Table 5:** Model parameters with their values. For interpretation, note that *α*_i_ parameters are within exponential functions. For instance, *α*_5_ = 0 corresponds to a minimum fecundity of exp(0) = 1.

| Parameter | Value | Interpretation |
| --- | --- | --- |
| $\alpha_1$ | 0.5 | juvenile predator survival – intercept |
| $\alpha_2$ | -0.01 | juvenile predator survival – slope |
| $\alpha_3$ | 0.5 | juvenile prey survival – intercept |
| $\alpha_4$ | -0.025 | juvenile prey survival – slope |
| $\alpha_5$ | 0 | predator fecundity – intercept |
| $\alpha_6$ | 0.004 | predator fecundity – slope |
| $\alpha_7$ | 2 | prey fecundity – intercept |
| $\alpha_8$ | -0.005 | prey fecundity – slope |
| $p$ | 0.7 | capture probability |
| $\phi_P^A$ | 0.7 | constant adult predator survival |
| $\phi_V^A$ | 0.6 | constant adult prey survival |
| $\sigma_y$ | 0.001 | observation error (negligible) |
| $n_P^A$ | 20 | initial number of adult predators |
| $n_P^J$ | 20 | initial number of juvenile predators |
| $n_V^A$ | 100 | initial number of adult prey |
| $n_V^J$ | 100 | initial number of juvenile prey |

## 4 Results

A typical simulation of the population counts (for one model run), together with their estimates under the fitted ICM (scenario 1) is presented in Fig. 2.

**Figure 2:**
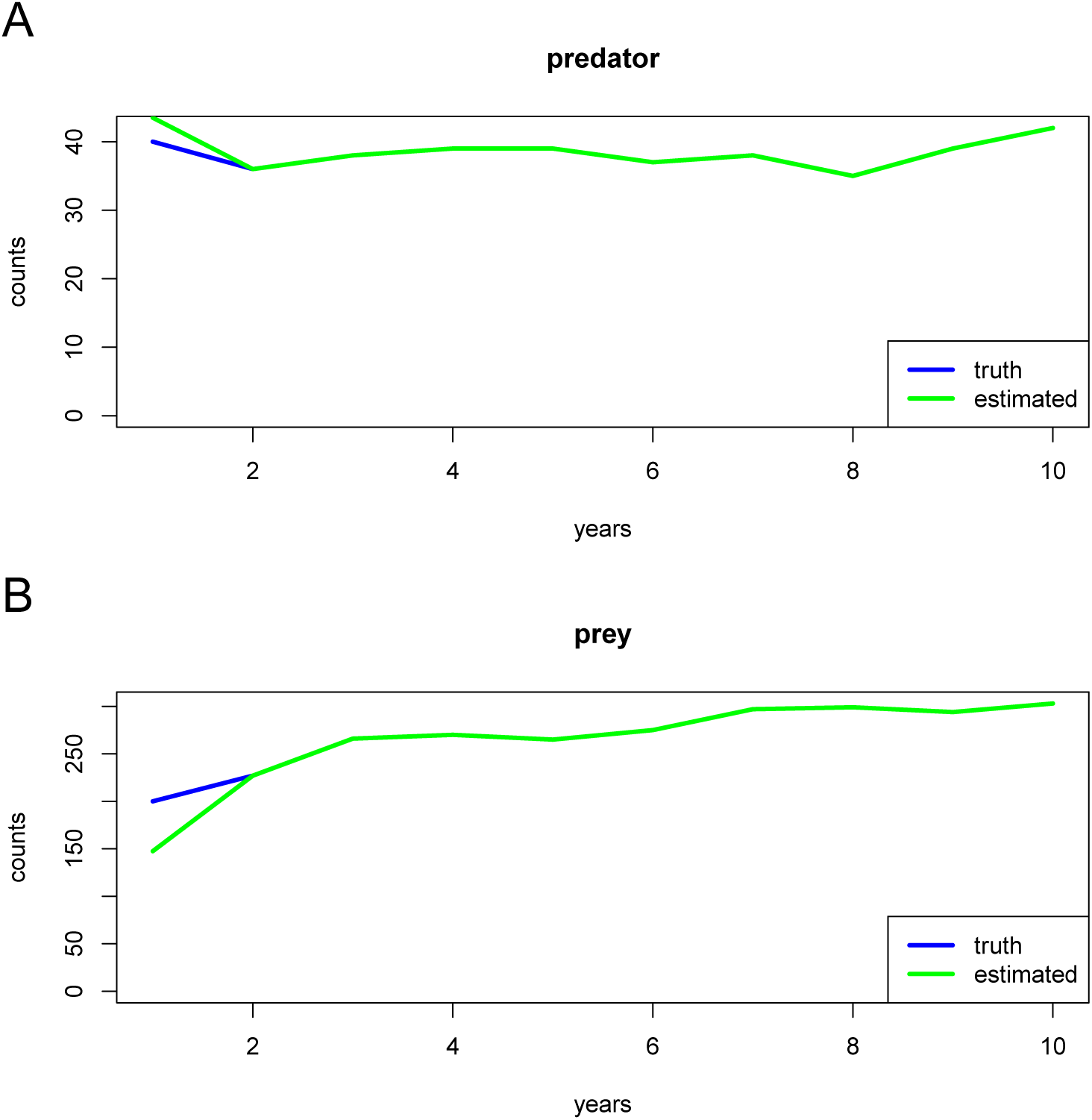
Time series of abundance (sum of both juvenile and adult stages) for predator (A) and prey (B)

We see that the predator and prey fitted trajectories match well the simulated trajectories. This is in fact true in all scenarios (not shown), and probably attributable to the high dimensionality of ICMs. During model fitting, the algorithm always finds a combination of vital rates that at least match the count trajectories. However, reproducing the shape of density-dependent relationships is a much harder task, as we show below.

In the scenario 1, the statistical fit does also reproduce well the four density-dependent vital rates (eq. 1, Fig. 3 show one typical statistical fit). This is true for most simulations in scenario 1. However, in the scenario 2, the fitted model could not reproduce density-dependencies in many cases. This bad match between simulated and fitted parameters in scenario 2 is made explicit by the examination of the bias and precision in *α*_*i*_ estimates (Fig. 4). Note that Fig. 4 does not present Bayesian credible intervals, though these exist as well (and we have examined them). In Fig. 4, we summarize the information provided by each of the 100 simulations by keeping the posterior mean of the parameter *α*_*i*_, which we denote 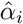. Presenting for 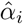 many simulations allows us to study the properties of this estimator of *α*_*i*_ in a similar manner to a frequentist study of the estimator. Bias is given by 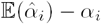 and precision by the dispersion (variance) of 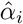.

**Figure 3:**
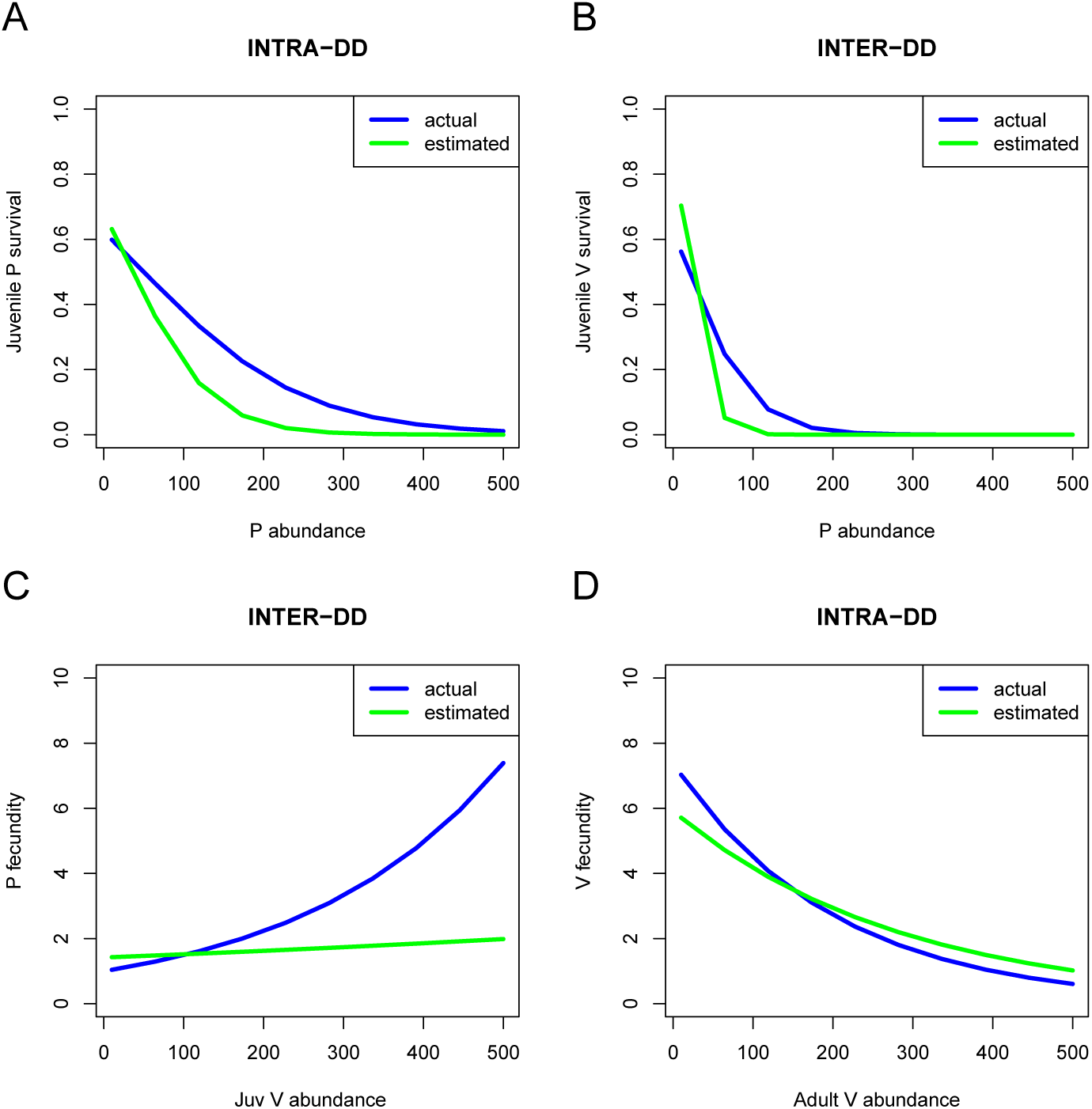
Density-dependencies for juvenile survival rates (A, predator and B, prey) as well as fecundities (C, predator and D, prey). Blue: simulated relationships, green: fitted relationships.

We see that the count-only scenario 2 generates large bias and low precision, in particular for interaction parameters (even indices for *α*_*i*_, *i* = 2, 4, 6, 8). Both scenarios 3 and 4 generate relatively good performances of the estimators for most *α*_*i*_. Interestingly, scenario 4 has almost identical performance to scenario 1 (almost the same precision, Fig. 4, and similar bias), which means that prey fecundity data is in fact of more value than prey capture-recapture data, at least for the parameters considered here.

To be thorough, we have considered our second parameter set with stronger predator-prey coupling (*α*_5_ = 0.5, *α*_6_ = 0.01, *α*_7_ = 1.5) and it shows similar results (Fig. B.1 in Appendix B). The slight systematic bias that exists for *α*_5_ in data scenarios 1, 3 and 4 vanishes for this other parameter set. This new parameter set also confirms the fact that scenario 4 is almost indistinguishable from scenario 1.

The same figure for the simulated *T* = 30 years observational study is presented in Appendix B (Fig. B.2). Combining data types still improves model performances: scenario 2 has the least precise estimates for most parameters. However, for *T* = 30, we observe less precise interaction estimates for scenarios 1, 3 and 4 (*α*_2_, *α*_4_) or with a little more bias (*α*_6_, *α*_8_) than for *T* = 10. The benefits of capture-recapture and fecundity data for the *T* = 30 setting, with a longer study but a less intense capture effort, are a little lower than for *T* = 10. This is logical and can be interpreted as a presence of proportionally more information in the counts for *T* = 30. The scenario 4 still brings low bias and estimates of similar quality to scenario 1, which confirms the good behaviour of this observational design for a different capture effort and time series length.

## 5 Discussion

Building on integrated population modelling, and combining it with nonlinear multispecies matrix models, we have developed a predator-prey Integrated Community Model (ICM). The density-dependent relationships in the ICM were designed to allow the model to persist for extended periods of time - this implied to not only consider density-dependent vital rates linking different species, but also density-dependencies within the same species. This resulted in four density-dependencies (Fig. 3). The model was structured and parameterized to correspond to a stage-structured predator-prey system composed of relatively large birds and/or mammals.

Using simulations of the ICM under contrasted data availability scenarios, we found that for a moderate study duration (10 years), the information contained within counts may be enough to reproduce the time series but not the density-dependencies. Inference about interactions therefore requires either longer time series (e.g. > 30 years or much more, which are often not available to practitioners) or a combination with other data types. The combination of all data types yielded estimates of vital rates and interaction parameters (density-dependent slope parameters) that had low bias and good precision, especially when compared to count-only estimates.

Although our results are promising, and definitely show that combining data sources can be worthwhile to estimate species joint dynamics, much remains to be done. We sketch below several areas for further research.

**Figure 4:**
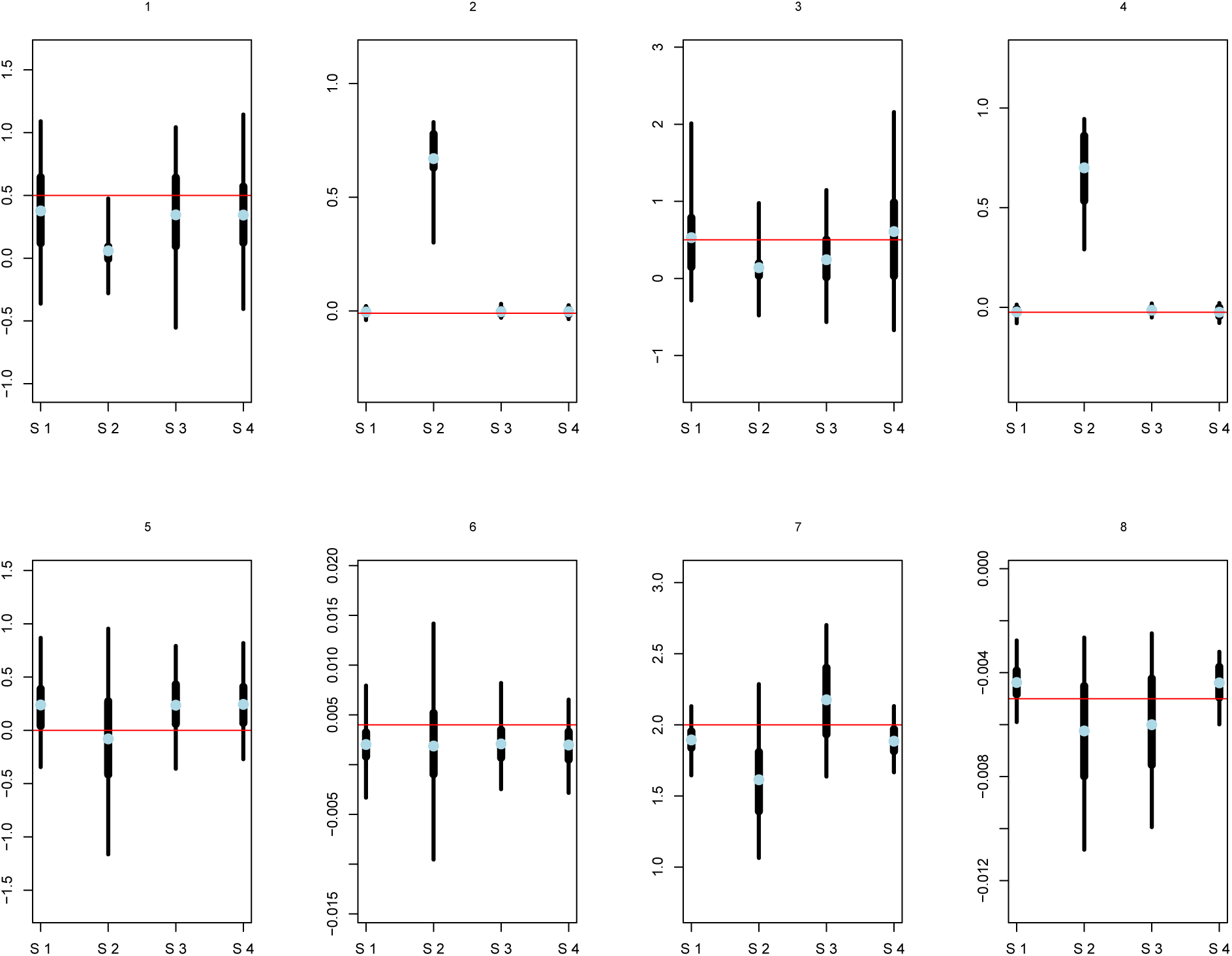
Bias and precision of *α*_*i*_ parameters, for *i* ∊ [|1 : 8|]. In each of the eights panels, four data scenarios (S 1 to 4) are considered. For each parameter and scenario, the thin bar represents a 95% confidence interval for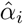; this interval is computed over 100 simulations, and thus quantifies the precision of the estimator. The wider bar is bounded by the 25% and 75% quantiles. The light blue dot represents the mean value 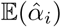 over 100 simulations; its distance to the true simulated parameter (red line) therefore quantifies estimator bias. Whenever the red line falls within the thin black bar, the CI contains the true parameter value so that the estimation is acceptable, otherwise it can be very poor (as in the count-only scenario 2).

### 5.1 Identifiability of the predator-prey ICM

We have shown that using all data types allowed estimating parameters with less bias and more precision. Weak identifiability *sensu* Gimenez *et al*. (2009) seemed approximately satisfied. An additional question is whether there are some redundant parameters in the model (Cole & McCrea, 2016). This issue can be explored to some degree using pairwise joint posterior plots (Appendix C, see Cole & McCrea 2016 for other ideas on how to pinpoint identifiability). Pair posterior densities show that parameters are (pairwise) uncorrelated unless they belong to the same density-dependency function. For instance, we see that *α*_1_ and *α*_2_ correlate negatively in the posterior distributions (Appendix C).

These negative correlations between the two parameters of a single density-dependency are a likely consequence of the parameterization of the model: the overall shape of the density-dependence curves can still remain unchanged using several parameter combinations. Whilst correlation between parameters, as well as ridges in the likelihood or joint posterior distribution are in many cases worrying, we show here that in this context, negative correlation in the posteriors for parameters of the same density-dependent curve is good rather than poor statistical behaviour: the correlation produces more precise estimated density-dependent relationships (Appendix C).

This could be explained by re-parameterizing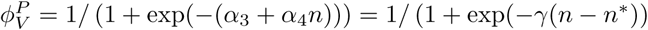, *n*^*\**^ being the equilibrium population size. We obtain *α*_3_ = *-γn*^*\**^ and *α*_4_ = *γ*, which produces a negative correlation. Assuming that *γ* is estimated less precisely than *n*^*\**^, we obtain the result. A similar argument for negative correlations in the posteriors can be made for the fecundity parameters: if *f*_*P*_ (*n*) = exp(*α*_5_ + *α*_6_*n*), then an increase of *δ* in *α*_5_ will produce a decrease of *nδ* if *f*_*P*_ is to stay approximately at the same value.

### 5.2 Refining the ICM

#### 5.2.1 Saturating functional forms for density-dependencies

In the current model, we consider a potentially unlimited increase in the predator fecundity (accelerating relationship). For the range of parameter values that we considered, this poses no problem because both prey and predator densities stay within reasonable bounds, but if the model is fitted or simulated in other contexts, this could become problematic: a surge in prey abundance would then allow for completely unrealistic predator fecundities. An avenue for improvement is therefore to implement a saturating predator fecundity, i.e.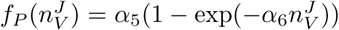. So far, all our attempts to do this ended with some notable bias on *α*_6_ (Appendix D), with the model estimating a flat rather than decelerating predator fecundity. Clearly more work in that area is needed.

#### 5.2.2 Observation error on the counts

In this article, our fitted models did not include observation error on the counts. We actually planned to do so, and our computer code allows this observation error to be implemented, but we found out through preliminary model fits that an unknown information error seems to induce major identifiability issues. The trajectories of predator and prey counts can still be reproduced when a poorly estimated observation error is present, but the density-dependent functions cannot. In other words, the model then produces good in-sample forecasts, but for the wrong reasons. All *α*_*i*_ values are estimated with strong bias in this case, including when all data types are used.

Our model belong to the general family of state-space models or SSMs (Newman *et al*., 2014) and as such, suffers from the same difficulties in implementing unknown observation errors in such models. Indeed, several modelling studies have reported that unknown observation errors in SSMs, in absence of multiple samples per time step (see in this case Dennis *et al*., 2010), can be quite problematic. Without prior information, observation error variances are often poorly identified, which then creates bias in other estimates (Knape, 2008; Lebreton & Gimenez, 2013; Auger-Méthé *et al*., 2016; Certain *et al*., 2018). Such biases are likely to be even stronger if the dynamics contains large fluctuations, especially chaotic ones, and more refined model fitting techniques may have to be employed (Wood, 2010; Hartig & Dormann, 2013).

Unless the observation error on the counts is known to some degree (e.g., because counts are estimated by a method reporting that error such as distance sampling) and can be specified as an informative prior, given the already high complexity of ICMs, we suggest to neglect it. However, if some degree of replication within a time step is available, it may be possible to estimate it (Dennis *et al*., 2010). We note in passing that although assuming an observation error on the counts is customary in population ecology – and we have followed this modelling tradition here – there is no special reason to think that counting animals is more subject to error than resighting or fecundity estimation (e.g., this may be true for ringed birds but not necessarily for mammals identified by fur patterns). Decision of including observation error on counts or not in a hierarchical model will therefore be best made on a case-by-case basis.

#### 5.2.3 Model structure

We have considered for now a pre-breeding census MPM for the ICM. However, we have also written out a post-breeding census version of the nonlinear matrix model, with a juvenile stage duration of potentially multiple years (Appendix E). Comparison of the timing of events implied by the two versions of the model (pre-versus post-breeding, e.g. Cooch *et al*. 2003) has revealed that only minor differences in interpretation are involved. This may be partly because our model, where vital rates are related to densities by nonlinear phenomenological functions, does not rely on explicit biomass conversion of prey into predators, i.e., litteral conversion of killed prey into newborn predators. Zhou *et al*. (2013) suggested that such discrete-time systems with biomass conversion might be more stable because of the ordering of events, assuming that predation occurs before reproduction. However, even though event ordering (reproduction vs survival) can affect stability in these models, census timing may not: Weide *et al*. (2018) show that, for models with explicit biomass conversion, census timing only imply minor quantitative differences in model outputs. For difference equations, major qualitative differences only occur between models that assume biomass conversion to happen before reproduction (more stable) or after (more oscillating), as shown by Weide *et al*. (2018).

A post-breeding census model brings our framework closer to a multi-species version of the density-dependent models of Neubert & Caswell (2000). This formulation, which we use for on-going theoretical work, was more internally consistent (Appendix E) but also more difficult to adapt directly in the IPM framework, which we adapted from previous equations and existing computer code for pre-breeding census models Abadi *et al*. (2012). On empirical grounds, both pre-breeding and post-breeding model can make sense depending on the timing of the census. If there are only two classes, juveniles and adults, in both the pre- and post-breeding census version, the corresponding individual-based community model can be described by a sequences of stochastic events involving only Poisson and Binomial distributions (Appendix E), Binomial for death rates and Poisson for recruitment. There is a good mapping between the ICM and the fully explicit stochastic IBM because the binomial sampling of a Poisson-distributed count still follows a Poisson distribution, and the binomial sampling of a Binomial-distributed count is still a Binomial distribution. However, real recruitment distributions might have added variance (e.g., Zero-Inflated Poisson or Negative Binomial) and these may be considered in future ICM work.

It is possible to extend the formulation of our model, so that juveniles are able to stay several years within the juvenile class (Appendix E). This may be a useful first step to model a longer-lived juvenile stage (i.e., more than a year). Indeed, using more than two classes requires to use multinomial distributions for transitions (Watkins, 2000) which amounts to specify the community-level IBM as a multi-type, density-dependent branching process (Haccou *et al*. 2005, instead of using Poisson and Binomial random variates as in section 2.2.1). This adds considerable complexity to the stochastic model and is left for further work.

Another layer of complexity that we have not tackled here is spatial structure. The spatial structure itself can contain information about interactions - although power can be an issue with spatial-only data (Rajala *et al*., 2018), spatio-temporal data could be of great use in helping to identify interactions. If only count data are spatial, counts might become increasingly important in identifying the model, potentially changing some of the results that we show here. Tredennick *et al*. (2017) show for instance that demographic data might not be needed to predict spatiotemporal, quadrat-level plant community dynamics. With that in mind, it is also important to realize that for many birds and mammals, count data are usually scarce and the spatial scales at which a count value can be defined can be very large, so that spatial replication in count data might be very hard to achieve. In some cases, it might even be easier to have spatially-dependent survival or recruitment data, since those can be individual-based (Chandler & Clark, 2014).

#### 5.2.4 More intricate or challenging dynamics

We have considered thus far that the predator-prey system exhibited a fixed point equilibrium (a consequence of slow bird and mammal life histories and predation on juveniles only), with added noise due to demographic stochasticity (section 2.2.1). There is therefore only a moderate ‘signal’ in the counts themselves. Changing parameters could allow for cycles or chaos to develop (Neubert & Caswell, 2000; Wikan, 2001), which could then provide more signal and make the count information more relevant, at least for long time series. Both Neubert & Caswell (2000) and Wikan (2001) have shown that these dynamics tend to occur for large fecundities: different systems than birds and mammals, involving for example fishes or plants might therefore have much wilder fluctuations. We confirm this for our model in Appendix A. However, a more fluctuating model can also be tricky to estimate because there can be multiple causation of oscillations (e.g., interactions of stochasticity and age structure, Greenman & Benton, 2004; Barraquand *et al*., 2017). It is therefore unclear what changing completely the range of parameters in this model – e.g., considering much faster life histories – would do to the quality of parameter estimates in the ICM.

Aside from changing parameters, one neat way to produce wilder dynamics would be to allow for predation on adults – this can be done phenomenologically rather than mechanistically by simply having an adult prey survival rate that depends on predator (adult) density. Other options are proposed by Zhou *et al*. (2013) and Weide *et al*. (2018). This would allow for predator-prey cycles (we have seen this in explorations of the nonlinear matrix model), but we note that we would need to change the time-frames of simulation considered (e.g., 30 to 100 years) in order to see well these long-period cycles. Such time-frames may be useful to model long-term ecological studies for which there is an historical time series of counts (say, 100 years) at the end of which we perform capture-recapture and fecundity estimation (during say, 10 or 20 years). This fits relatively well the data that could be obtained on lynx and hare (Krebs *et al*., 2018).

Another worthy exploration would be interaction estimates when there are actually no true linkages between the species’ population dynamics (i.e., no interaction at all, not even unmodelled indirect interactions). In the present article, we asked how different data combinations may or may not allow the model to recover interactions that are known to be present. This fits well predator-prey pairs for which there is prior knowledge about potential impacts of predators on their prey, which is clearly the situation we placed ourselves in. However, for less well-understood ecological configurations, there is always a risk of error in assessing who interacts with whom, and the question of what the ICM would infer if two species have in fact independent dynamics may be paramount. We think that such cases would be especially relevant when predator diet is poorly known or in a competition context, as it can be difficult to evaluate whether or not two species truly share a similar niche.

### 5.3 Suggestions for empirical surveys aiming at parameterizing multi-species models

We would like to finally conclude on what data should be collected in the field, if the aim is to parameterize multi-species and stage-structured dynamic models. Our results show first that longer time frames are not necessarily best (assuming that there is a trade-off between study duration and capture effort per year), especially for vertebrates. This is because 30 years of counts alone contain limited information to parameterize high-dimensional models (Barraquand & Nielsen, 2018), and capture-recapture and reproduction data add much needed independent information on parameters. Moreover, for the parameter values that we considered, tuned to birds and mammals, a prey fecundity survey brought more precision to interaction and vital rate estimates than a capture-recapture survey. Top predators are usually well-studied, notably due to conservation concerns, but their more abundant prey could be difficult/costly to follow through capture-recapture and be simply counted. Our results suggests that a prey fecundity survey adds much value, probably at a small fraction of the cost of a capture-recapture program.

In a Bayesian framework, a priori knowledge on some of those parameters can also be gained from the MPM databases (Salguero-Gómez *et al*., 2016), which will be especially important for poorly studied species. Which parameter(s) should be set with prior information will then depend on a trade-off between the degree of phylogenetic conservatism of the parameter (Che-Castaldo *et al*., 2018) and the ease which with the information can be collected in the field. Adult survival is a good target (costly to infer, probably fairly conserved between related species), and the analyses that we performed could be attempted again using a mixture of real data and database-derived a priori information.

## Acknowledgements

OG was funded by the French National Research Agency (grant ANR-16-CE02-0007) and FB by LabEx COTE (ANR-10-LABX-45). We thank Bachar Tarraf for discussions on nonlinear matrix models, as well as David N. Koons and an anonymous referee for thoughtful remarks and references.

## Author contributions

OG and FB developed the project. FB constructed a first version of the predator-prey model and OG a first version of code integrating datasets. Both authors then contributed to subsequent versions of models and code. The article contents and structure were jointly decided. FB wrote a first draft, which was then edited by both authors.

## Code availability

The computer code for simulating and estimating the model can be found in the following GitHub repository https://github.com/oliviergimenez/predator_prey_icm [This repository will be made public upon acceptance and deposited into Zenodo to have an associated DOI].

## Appendices

### A Trajectories of deterministic and stochastic versions of the nonlinear MPM

We present in Fig. A.1 simulations showing the convergence of the model to a fixed point for our first parameter set (Table 5 in main text) and second parameter set (with changed parameters *α*_5_ = 0.5, *α*_6_ = 0.01, *α*_7_ = 1.5, see also Appendix B). Demographic stochasticity does generate more or less fluctuations around this fixed point though (Fig. A.1(b) and (d)).

**Figure A.1:**
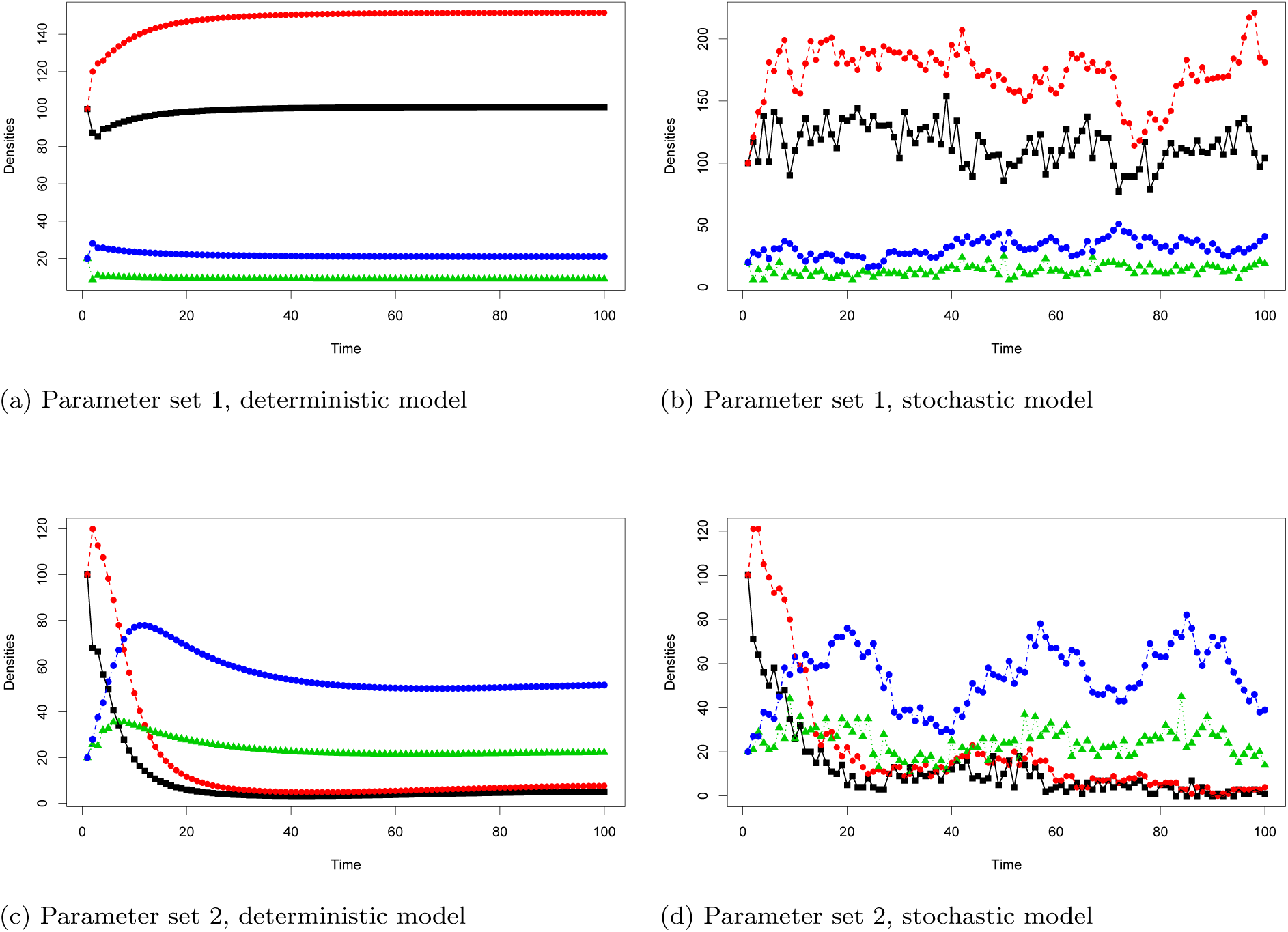
Comparison of deterministic and stochastic model trajectories for two parameter sets. Black squares, juvenile prey; Red filled circles, adult prey; Green triangles, juvenile predator; Blue filled circles, adult predator.

The stability of this model (in the Lyapunov sense, cf. Kot (2001) for example) is lost when intra-specific density-dependencies are removed (Fig. A.2), where trajectories show diverging oscillations.

**Figure A.2:**
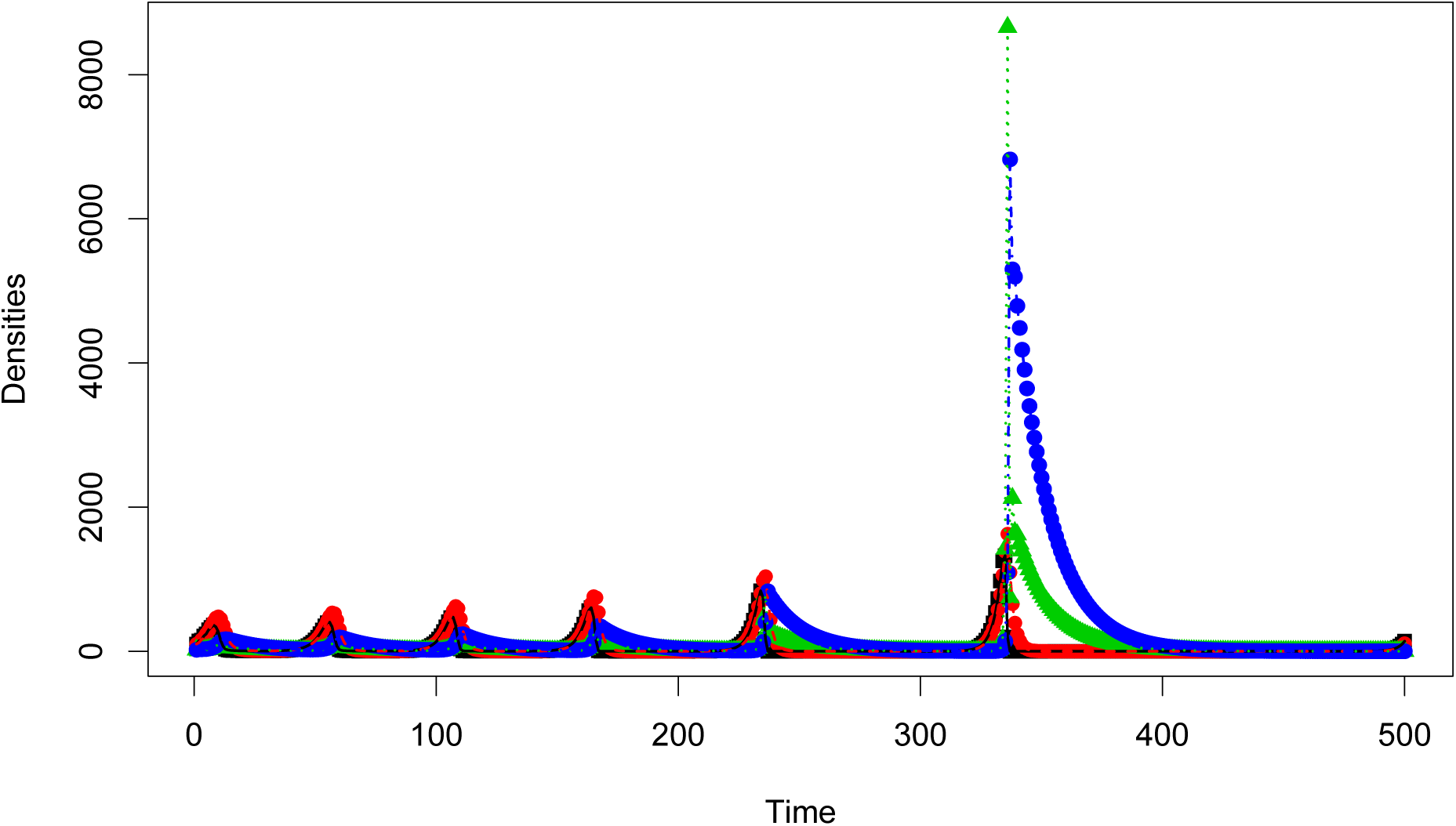
Diverging oscillations of the model with interspecific interactions but without intraspecific interactions.

We now present different dynamics obtained by increasing maximal fecundity to exp(5) for parameter set 1 and exp(4.5) for parameter 2 in Fig A.3. The model produces oscillations (likely invariant loops or chaos, more detailed investigations would be needed to distinguish between those). For a mathematical investigation of similar dynamics see for instance Wikan (2001).

The stochastic versions are also oscillating (Fig. A.3(b)), unless the high predator-prey coupling generates extinction when combined to the demographic stochasticity (small prey numbers and sampling create extinction, Fig. A.3 (d)).

**Figure A.3:**
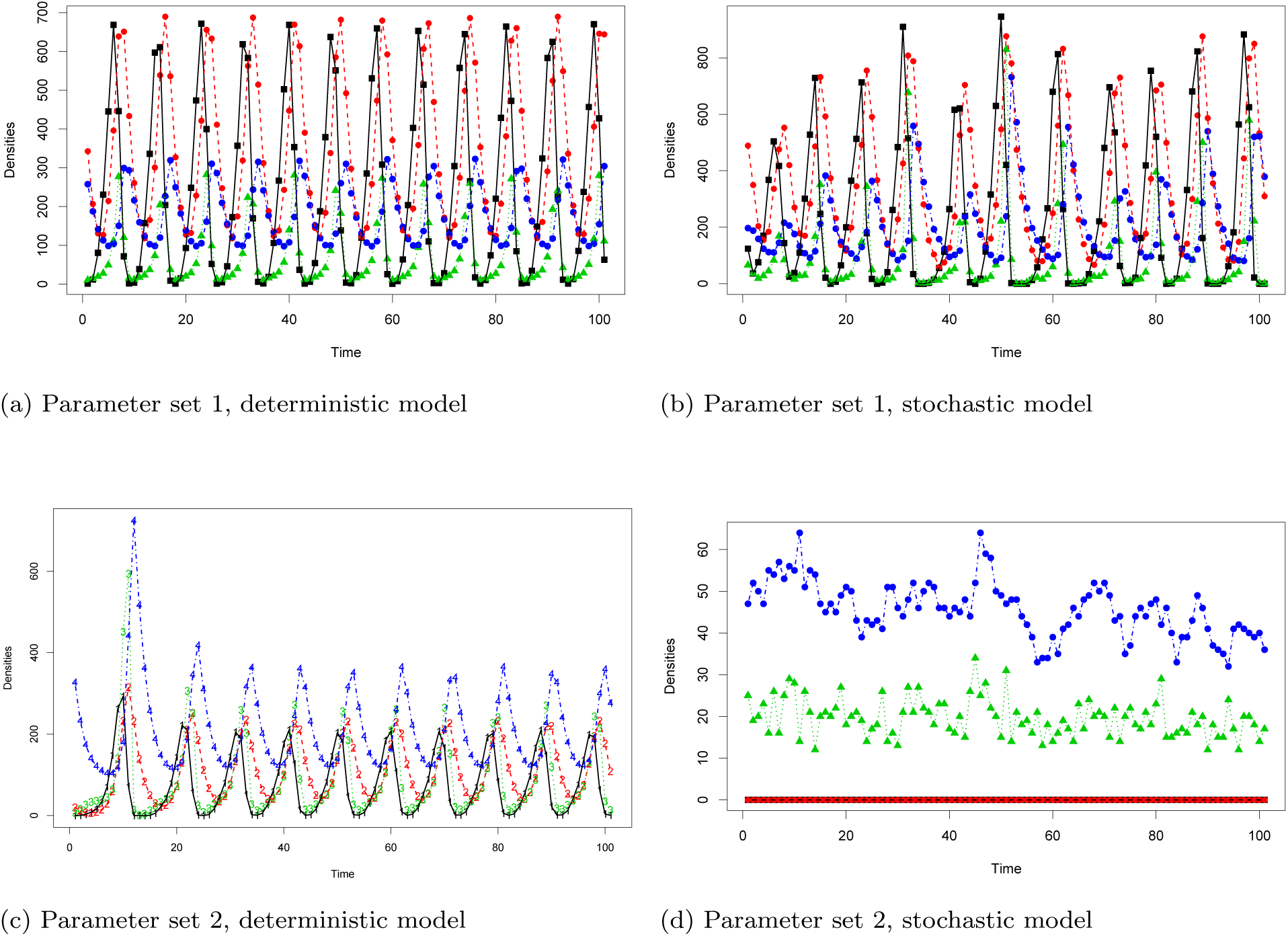
Comparison of deterministic and stochastic model trajectories for two parameter sets.

### B Additional estimates of alpha parameters

#### Results for the second parameter set

The second parameter set that we considered differed from the first by the following: *α*_5_ = 0.5, *α*_6_ = 0.01, *α*_7_ = 1.5, which generated a stronger predator-prey coupling.

**Figure B.1:**
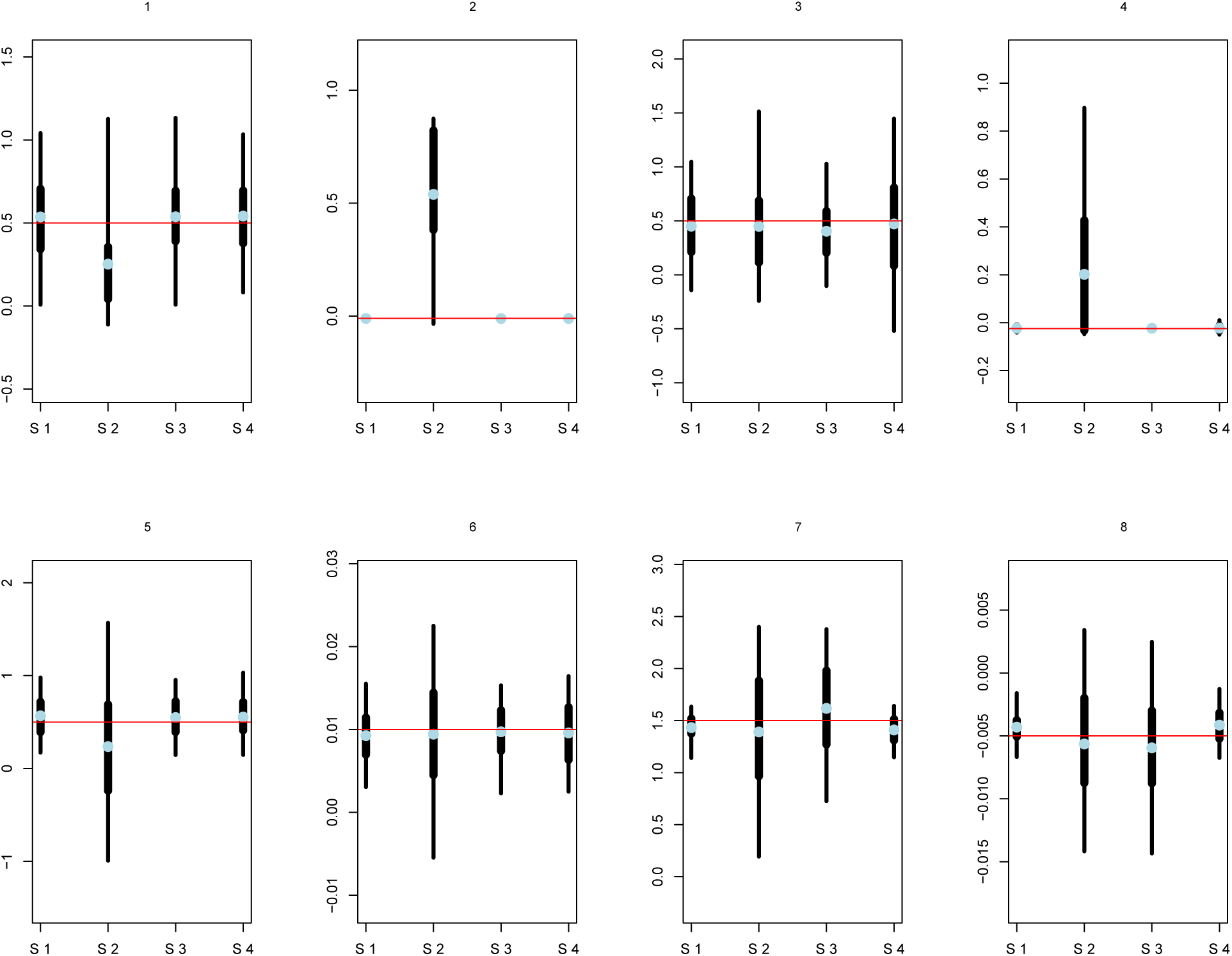
Bias and precision of α_i_ parameters, for *i* ∊ [|1 : 8|]. In each of the eights panels, four data scenarios (S 1 to 4) are considered. For each parameter and scenario, the thin bar represents a 95% confidence interval for 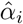; this interval is computed over 100 simulations, and thus quantifies the precision of the estimator. The wider bar is bounded by the 25% and 75% quantiles. The light blue dot represents the mean value 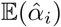 over 100 simulations; its distance to the true simulated parameter (red line) therefore quantifies estimator bias.

##### Results for the *T* = 30 years observational design

**Figure B.2:**
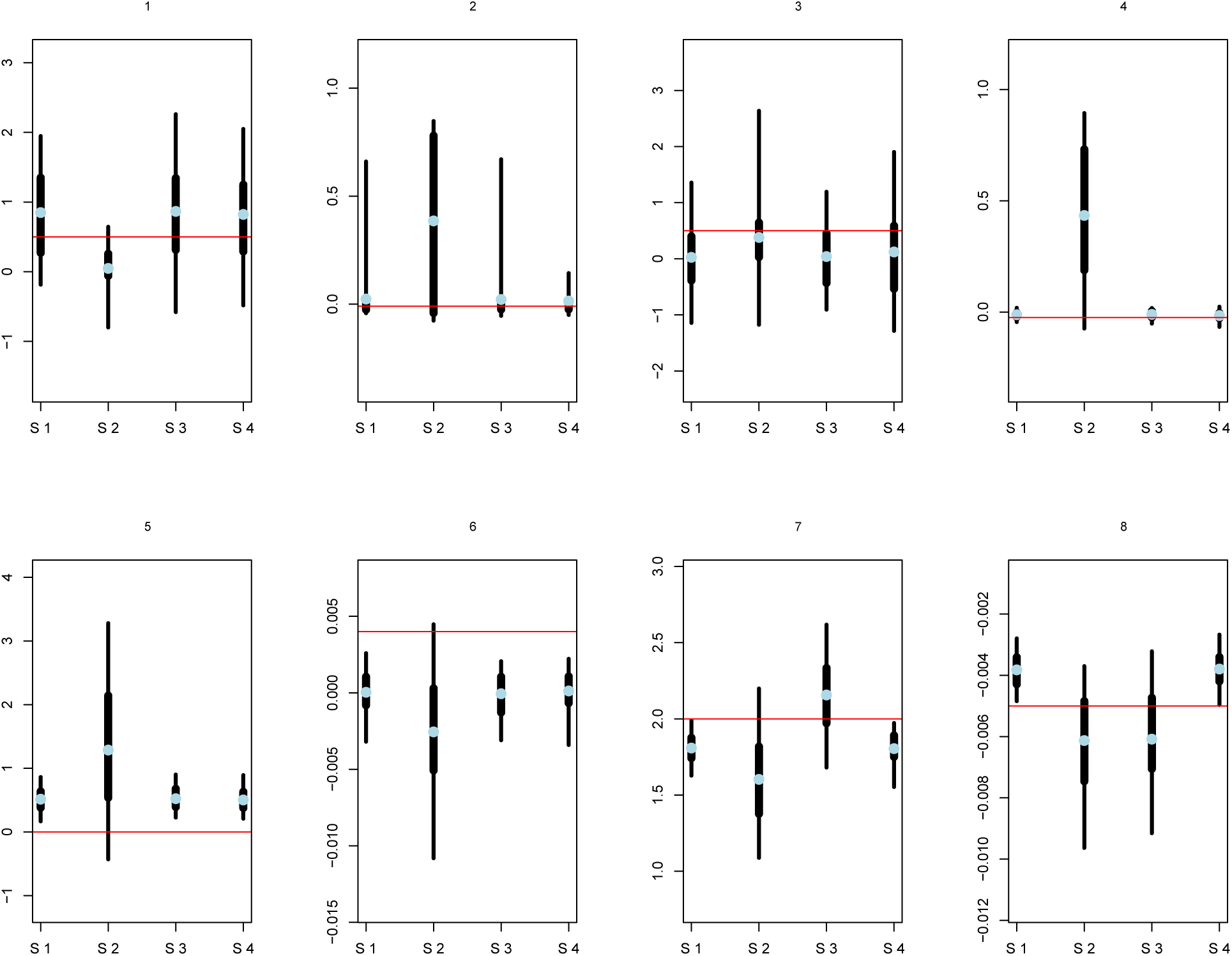
Bias and precision of α_i_ parameters, for *i* ∊ [|1 : 8|]. In each of the eights panels, four data scenarios (S 1 to 4) are considered. For each parameter and scenario, the thin bar represents a 95% confidence interval for 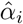; this interval is computed over 100 simulations, and thus quantifies the precision of the estimator. The wider bar is bounded by the 25% and 75% quantiles. The light blue dot represents the mean value 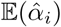 over 100 simulations; its distance to the true simulated parameter (red line) therefore quantifies estimator bias.

### C Identifiability diagnostics

The following plots describe the joint posterior distribution for pairs of parameters (based on two MCMC chains). Considerable linkage between the parameters that belong to the same density-dependent relationship are visible, these manifest as an elongated ellipsoid instead of a circle-shaped joint posterior for the pair. We present these diagnostic plots for *T* = 10 (Fig. C.1, scenario 1; Fig. C.2, scenario 2) and *T* = 30 (Fig. C.3, scenario 1; Fig. C.4, scenario 2).

For *T* = 10, we have good convergence (Gelman and Rubin’s *R <* 1.1); for *T* = 30, convergence and chain mixing are less good, which generates some bimodality on these posterior graphs.

**Figure C.1:**
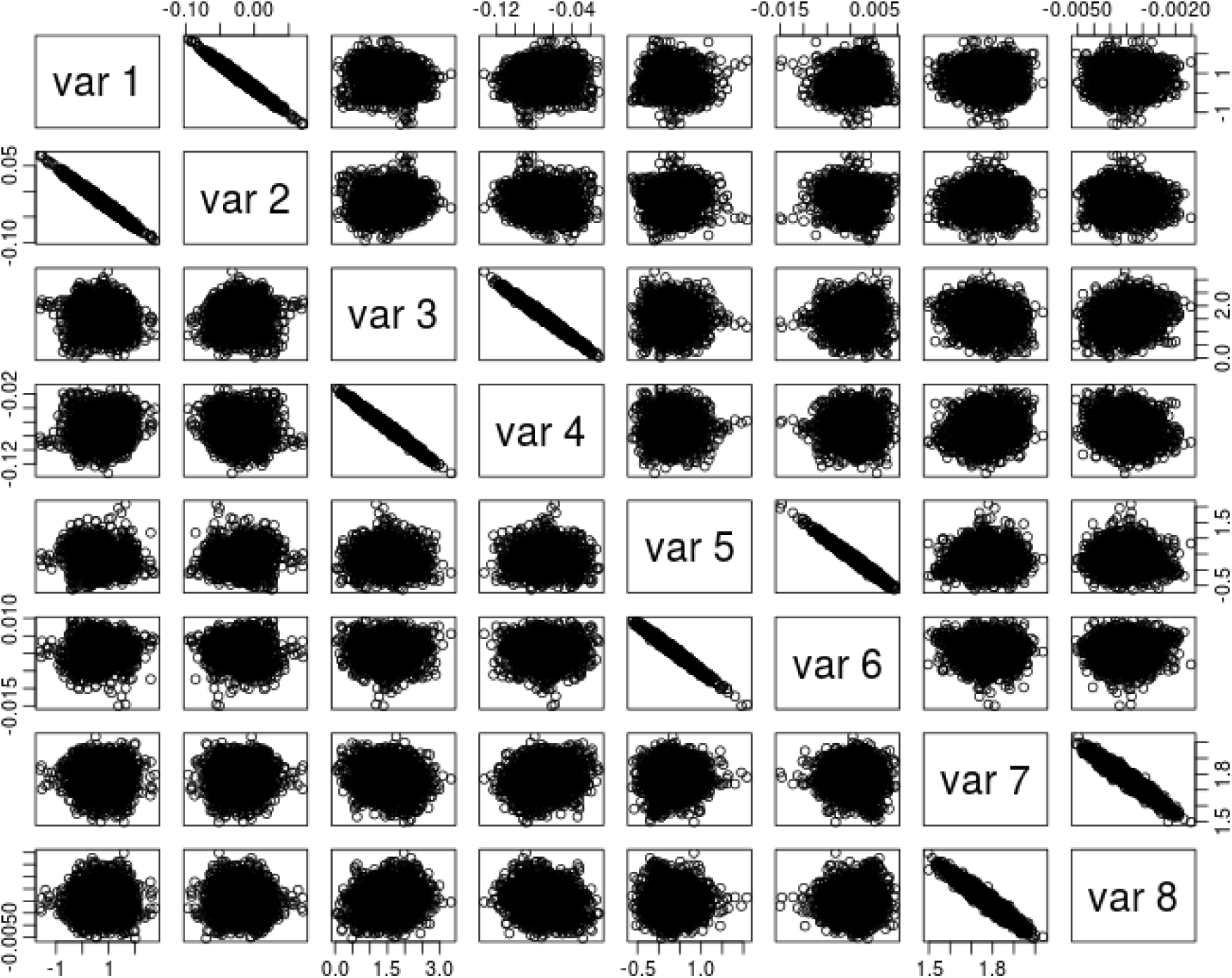
Pair posterior distribution for *α*_*i*_ parameters.

**Figure C.2:**
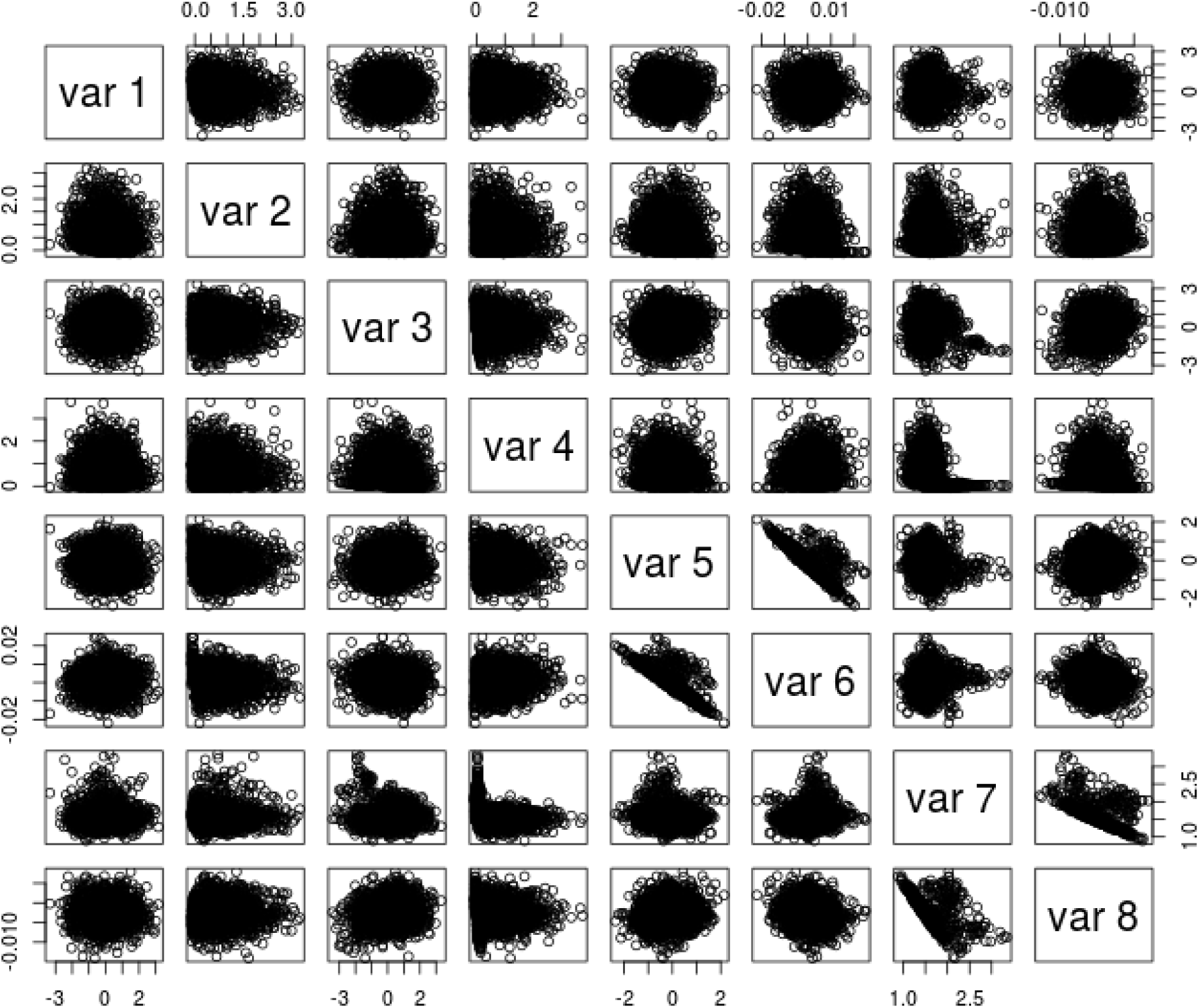
Pair posterior distribution for *α*_*i*_ parameters.

**Figure C.3:**
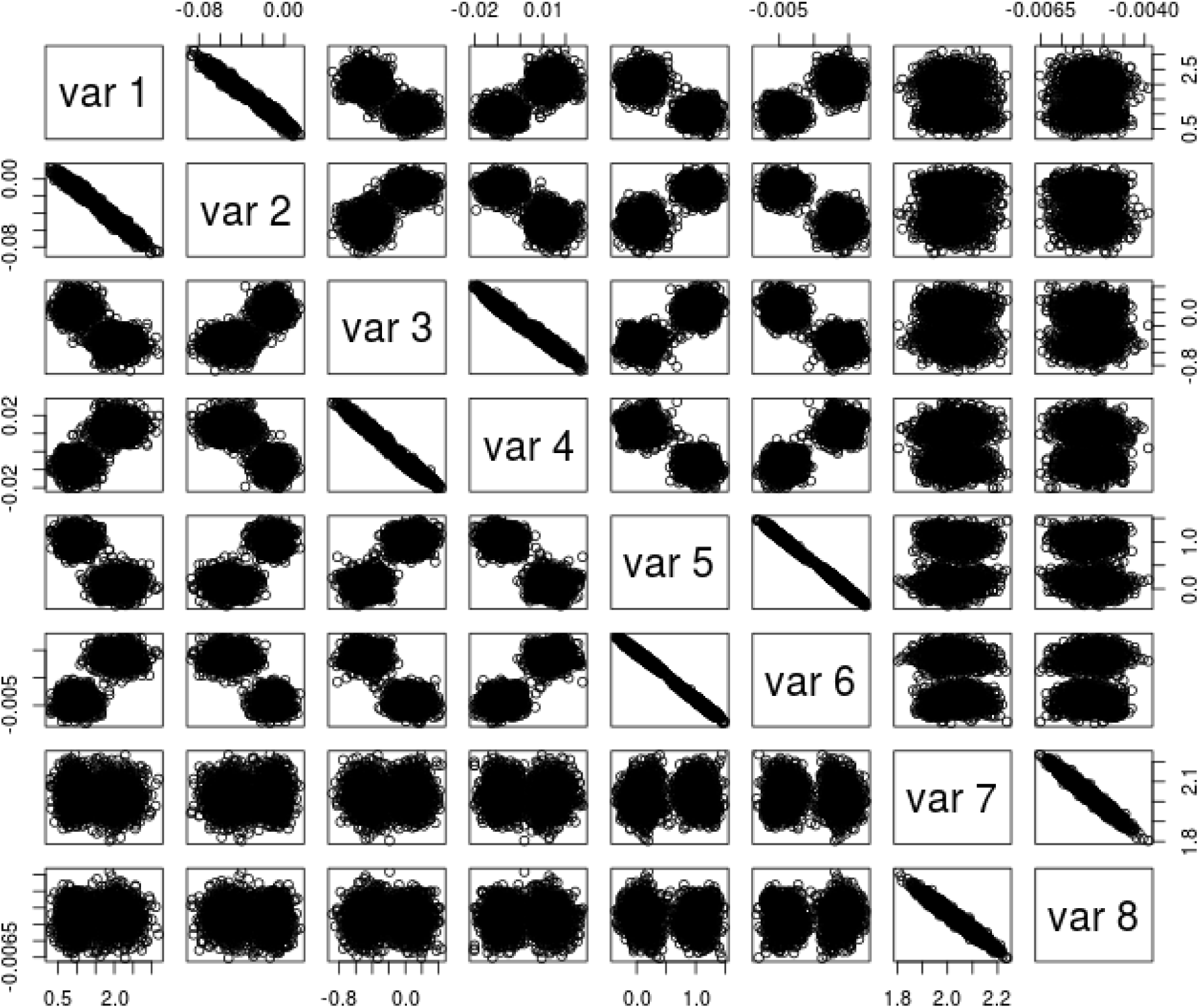
Pair posterior distribution for *α*_*i*_ parameters.

**Figure C.4:**
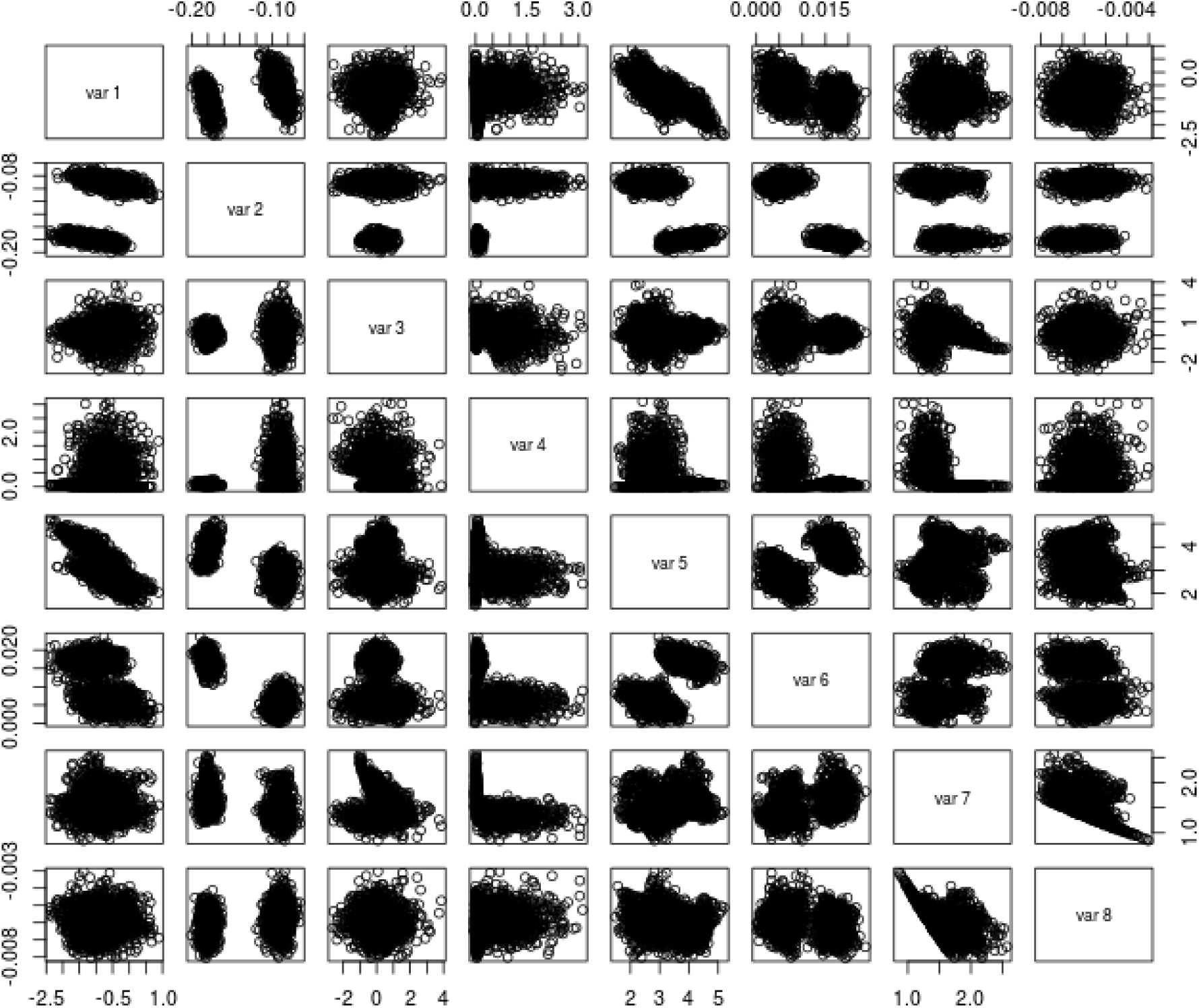
Pair posterior distribution for *α*_*i*_ parameters.

The persistent correlation in the posterior distribution of (*α*_1_, *α*_2_), (*α*_3_, *α*_4_), etc., for scenario 1 can be shown to have very little detrimental effect on the shape of the curves (Fig. C.5). The correlation between parameters even contributes to increase the precision of the curve, since randomizing (shuffling both vectors for *α*_3_ and *α*_4_ independently) decreases the precision of the curve (Fig. C.6). In Fig. C.6, the inflection point of the curve is in fact less marked than in Fig. C.5 that incorporates the correlation in the joint posterior.

**Figure C.5:**
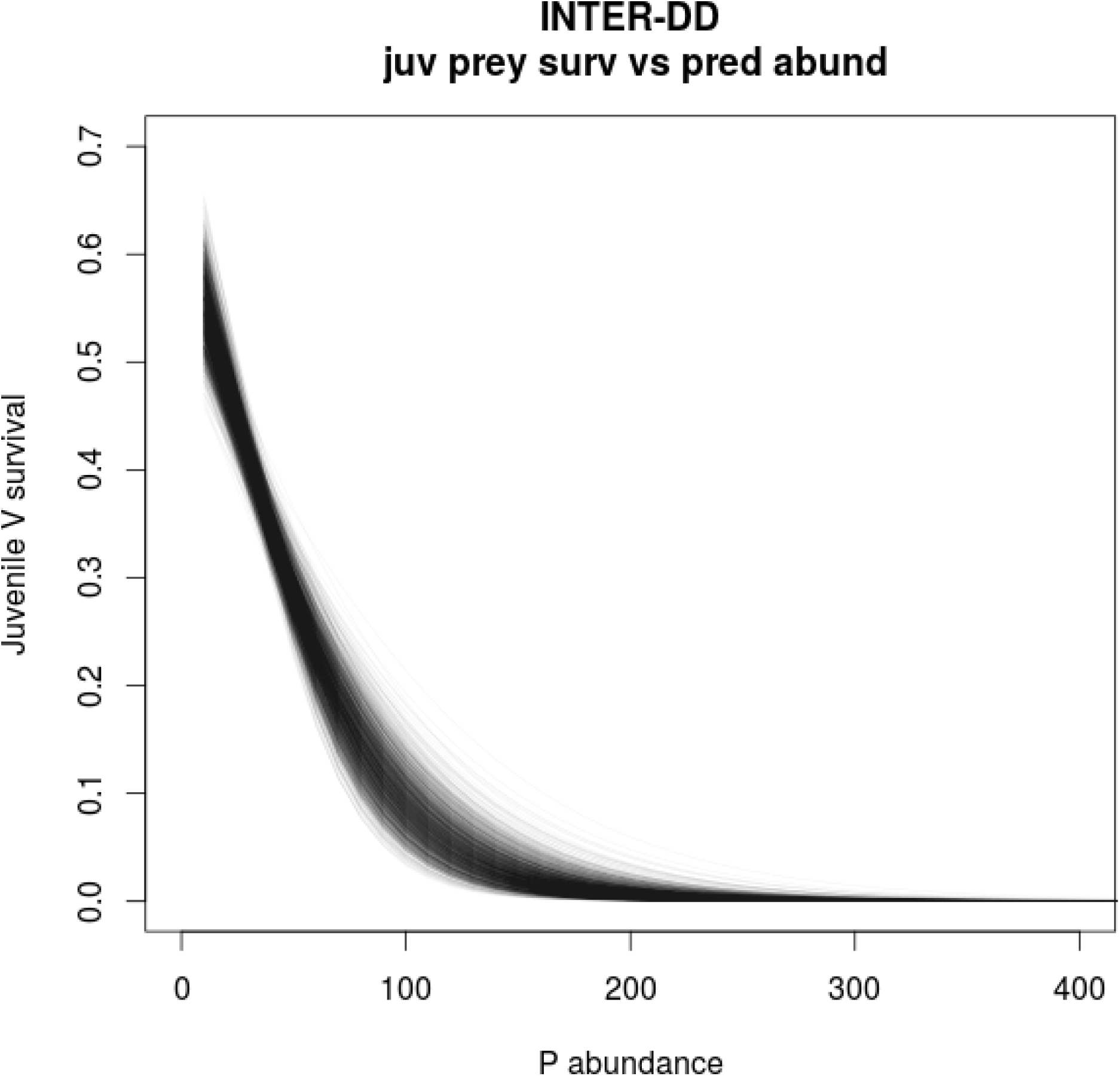
Prey juvenile survival curve including the correlation between *α*_3_ and *α*_4_. The plot has been constructed by using all the (*α*_3_, *α*_4_) values of the joint posterior distribution. The shading indicates the most probable values.

**Figure C.6:**
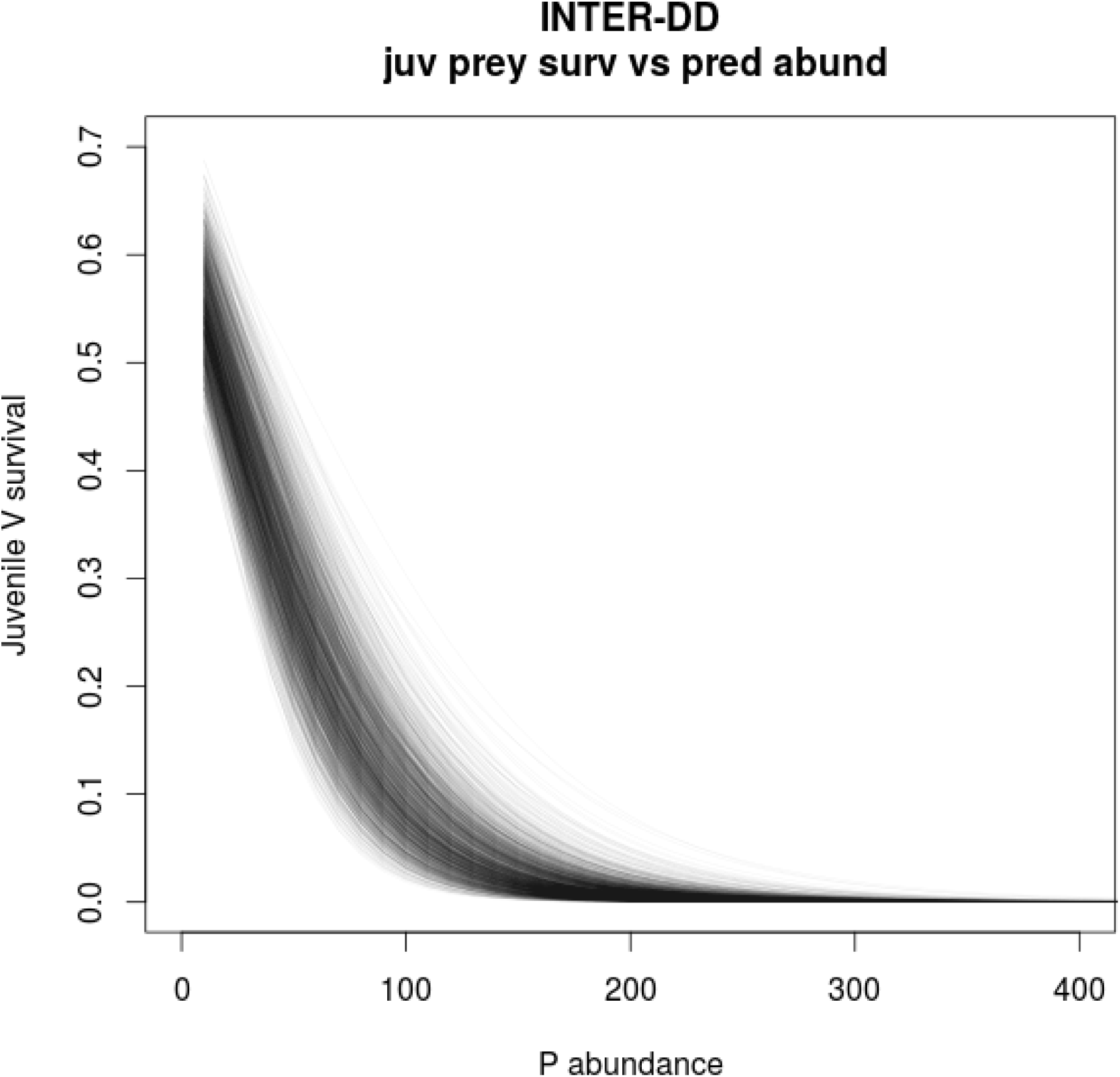
Prey juvenile survival curve without the correlation between *α*_3_ and *α*_4_. The plot has been constructed by shuffling independently *α*_3_ and *α*_4_ values. The shading indicates the most probable values.

### D Nonlinear predator fecundity

We considered both a saturating negative exponential function (Fig. D.1, 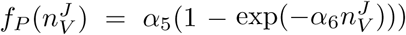 and a Michaelis-Menten function (Fig. D.2, 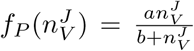. More informative priors improved the fit for other relationships (Figs. D.3,D.3) but did not correct the problem of ‘flat’ estimation of predator fecundity.

**Figure D.1:**
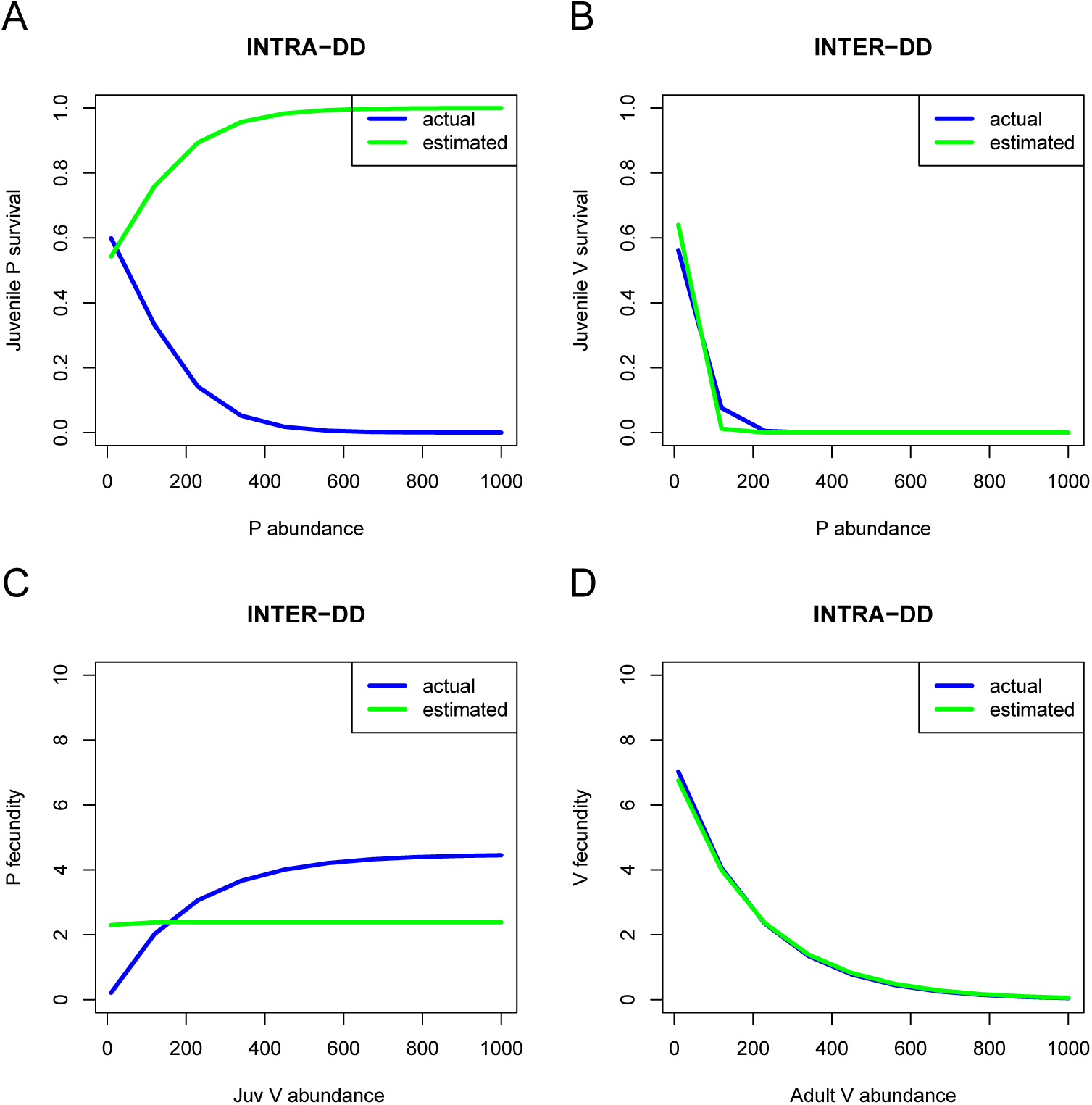
Saturating negative exponential predator fecundity. Density-dependencies for juvenile survival rates (A, predator and B, prey) as well as fecundities (C, predator and D, prey). Blue: simulated relationships, green: fitted relationships.

**Figure D.2:**
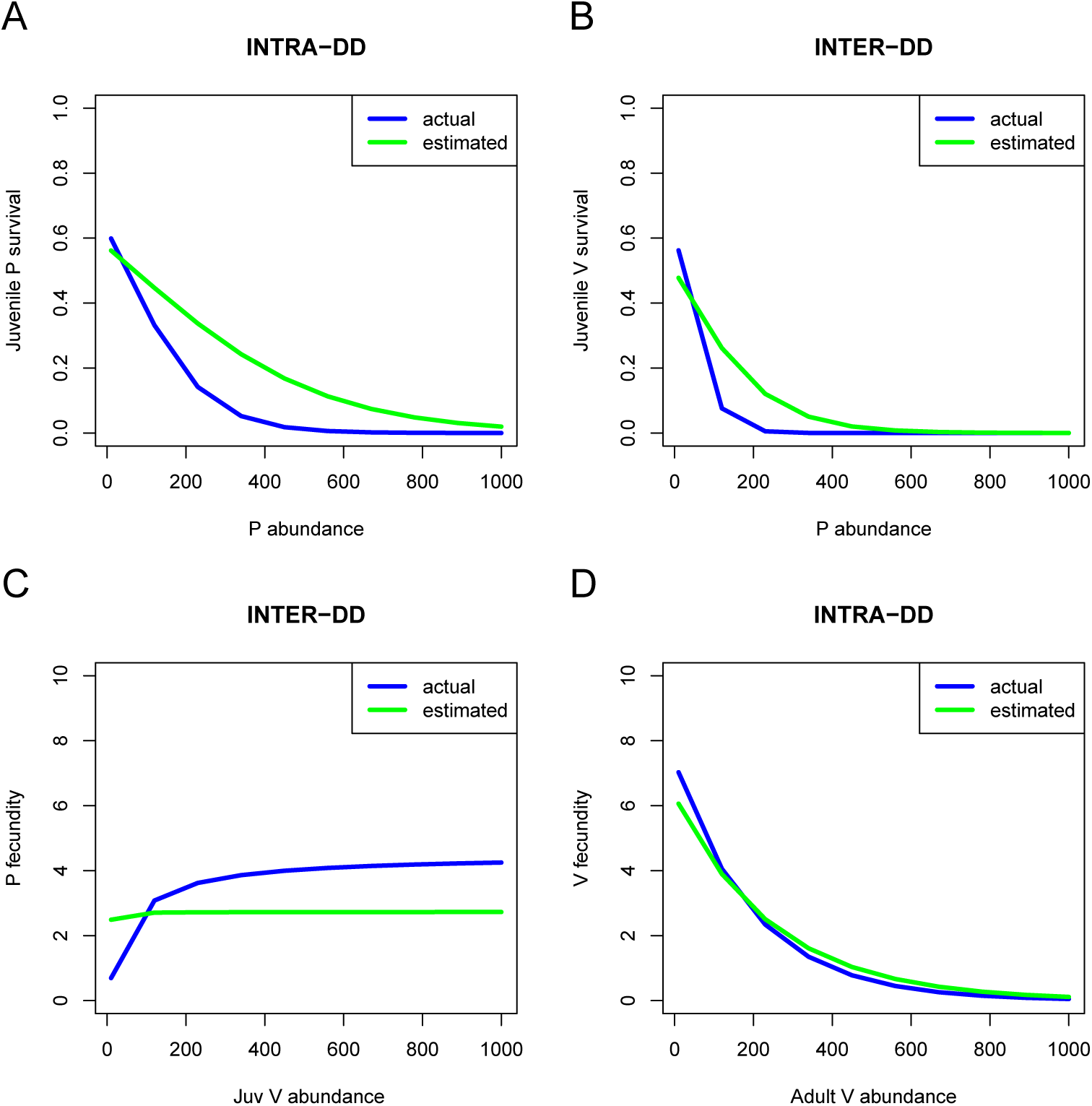
Saturating Michaelis-Menten predator fecundity. Density-dependencies for juvenile survival rates (A, predator and B, prey) as well as fecundities (C, predator and D, prey). Blue: simulated relationships, green: fitted relationships.

**Figure D.3:**
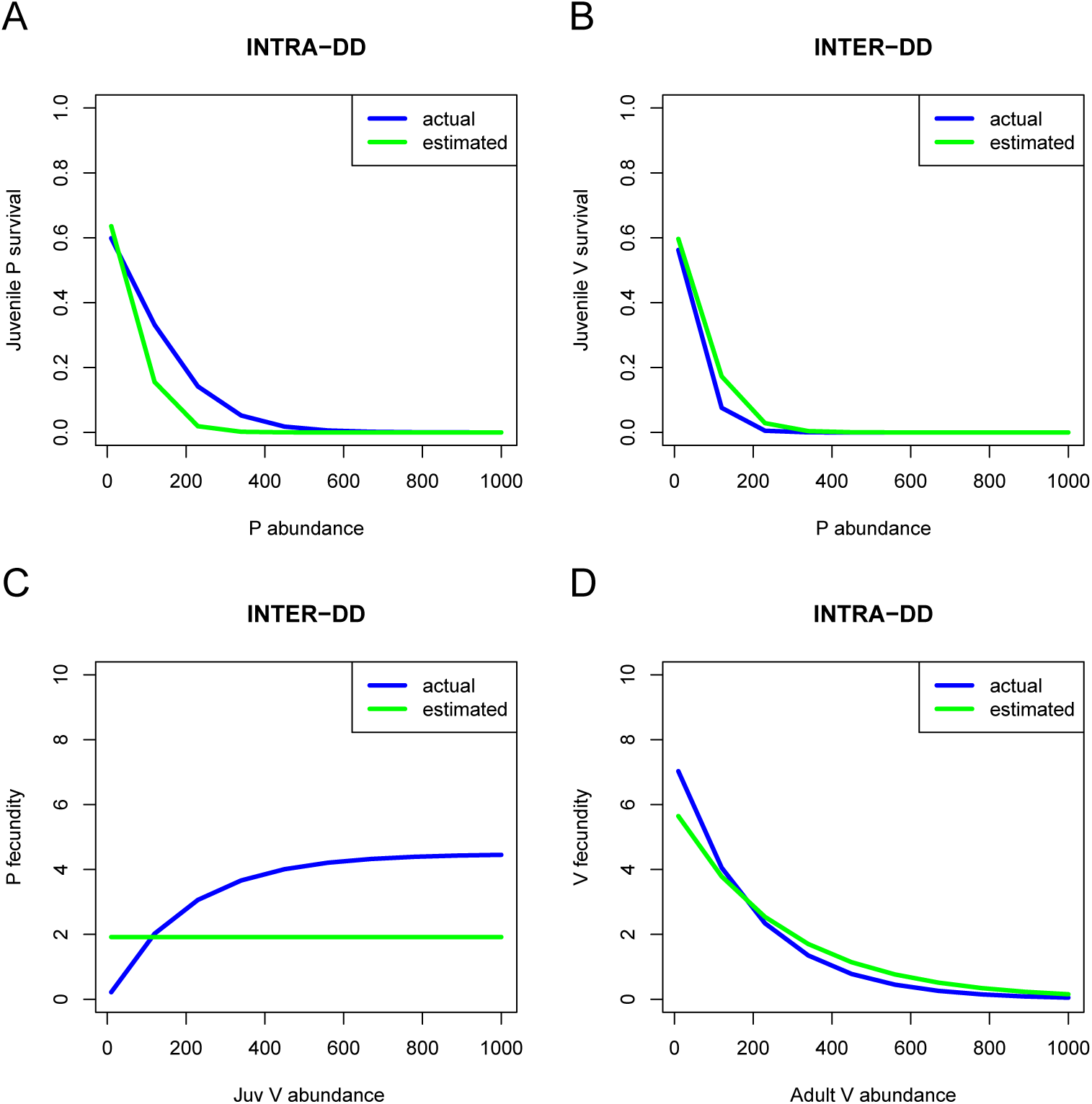
Saturating negative exponential predator fecundity and more informative priors. Density-dependencies for juvenile survival rates (A, predator and B, prey) as well as fecundities (C, predator and D, prey). Blue: simulated relationships, green: fitted relationships.

**Figure D.4:**
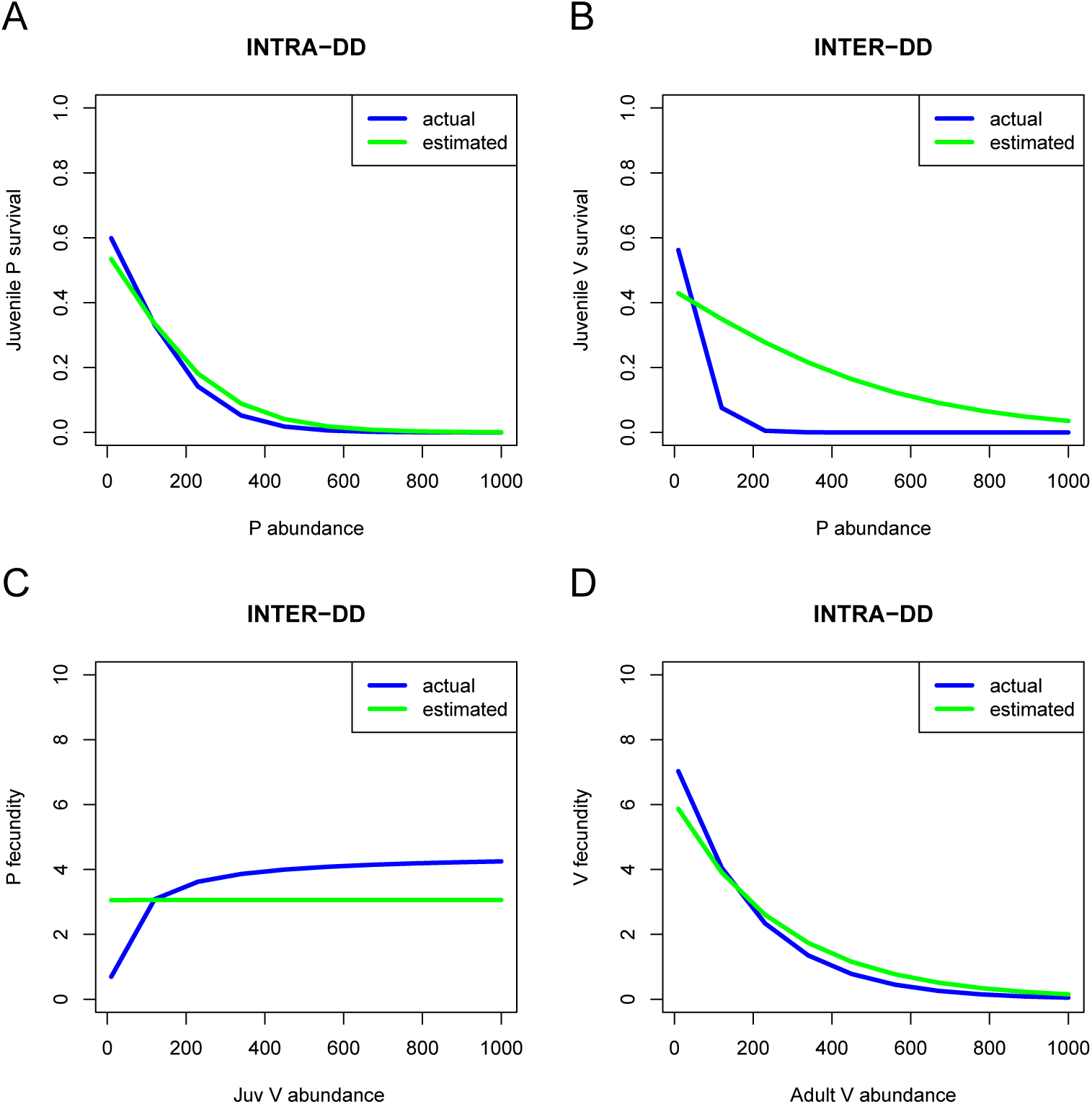
Saturating negative exponential predator fecundity and more informative priors. Density-dependencies for juvenile survival rates (A, predator and B, prey) as well as fecundities (C, predator and D, prey). Blue: simulated relationships, green: fitted relationships.

### E Pre- and post-breeding census model for *γ <* 1

#### E.1 Pre- and post-breeding matrix model for *γ <* 1

The general equation of our nonlinear matrix model is

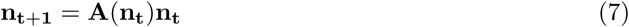

for the counts of the two species and two stages per species, with

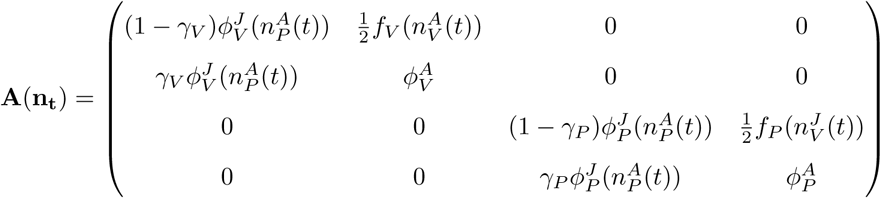

in the post-breeding case (similar to Neubert & Caswell 2000) and in the pre-breeding case (this article),

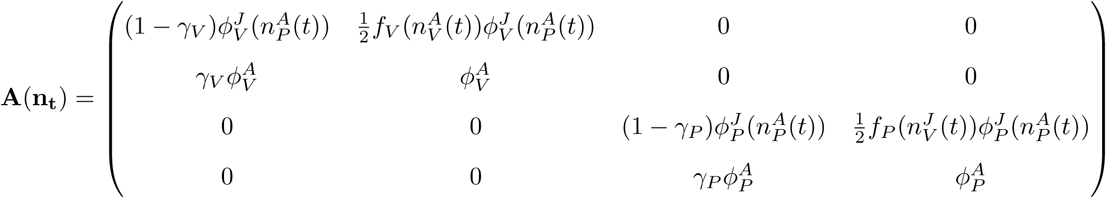

All other notations are similar to the main text. These two formulations with different census dates lead, interestingly, to slightly different versions of the corresponding stochastic individual-based model (IBM). We provide below a full derivation for the IBM.

#### E.2 Order of events in the Stochastic IBM

All events can be ordered on a time arrow by introducing an intermediate time *t ′* between *t* and *t* + 1 (see also Cooch *et al*. 2003; Weide *et al*. 2018). We consider first the post-breeding census scenario.

#### Post-breeding census

We assume that before census, reproduction occurs at some time *t ′* reasonably close to *t* + 1. At time *t* starts the survival phase, for both stages. We will derive the IBM for the prey population dynamics.

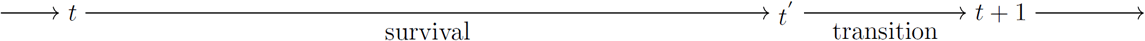

We have the update equation for juvenile prey population size at time *t* + 1,

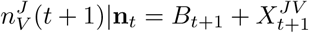

where 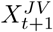 is the number of juveniles born in the previous time period that survive up to *t* + 1.

It therefore includes all the juveniles from the previous time period that

1. have survived (survival happens first)
2. have not transitioned to adulthood (transition happens second).

Because *t′* is very close to *t* + 1 (post-breeding census), we can neglect the survival fraction of juveniles of the year so that *B*_*t*+1_ = *B*_*t*_′ and

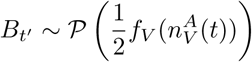

On the other hand, we have the number of individuals staying juveniles after *t′*

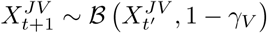

and finally the number of survivors from 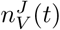

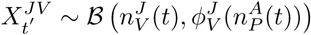

so that

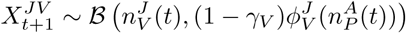

This follows from the fact that if *X|Y ∼ ℬ* (*Y, p*) and *Y ∼ ℬ* (*n, q*) then *X ∼ ℬ* (*n, pq*).

Let us denote 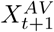 the individuals that have matured to adulthood between *t′* and *t* + 1, then 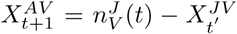, as we substract from the previous population size of juveniles individuals that have not maturated^1^.

The number of adults at the next iteration is then obtained as

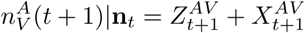

Where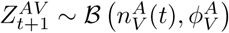. The prey part of the stochastic IBM is now fully specified.

Similar arguments follow for the predator compartment. The time ordering in this model means that from the prey point of view, predation happens on juveniles born in the previous year (if *γ* = 1, these would be between a few months and a little more than a year old). Predation occurs continuously during the (long) survival phase. Birth is always, in such discrete-time models, a pulse event (i.e., occurring at once or almost) at a given time *t′*.

For the predator population dynamics, predator reproduction – which happens at *t*′ as well – is driven by the abundance of the juvenile *at the beginning of the survival process starting at t*. Predator reproduction is therefore very much influenced by the prey individuals born last year. The prey and predator parts of the model are therefore consistent.

#### Pre-breeding census

Now we consider a pre-breeding census for which the time arrow is slightly differently organised, with birth at *t′* right after census at *t*.

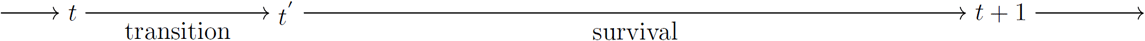

As in the previous derivation, we consider that the juvenile population size at time *t* + 1 is given by the following update equation

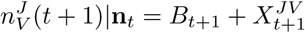

This translates the fact that juveniles are composed of both newborns and surviving juveniles that did not mature between *t* and *t* + 1. The difference now, for the pre-breeding census, is that there is a long period during which the juveniles of the year need to survive so that *B*_*t*+1_ ≠ *B*_*t*_′. In fact,

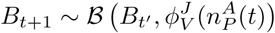

with a production of chicks at *t′*

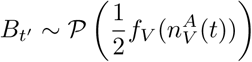

This leads to a marginal distribution of *B*_*t*+1_

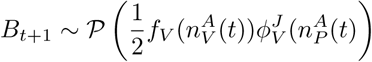

Indeed, if *X∣Y ∼ ℬ* (*n, p*) and *Y ∼ 𝒫* (*λ*), then *X ∼ 𝒫* (*pλ*). Here, predation – modelled through the predator-dependence of juvenile survival – occurs as well on *juveniles of the year*^2^, so eggs or young chicks. The surviving juveniles from the previous year are given by

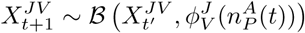

themselves picked from the individuals that did not mature

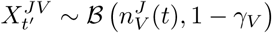

so that

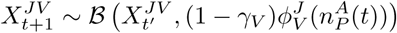

For the adults,

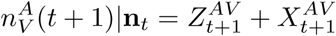

with surviving old adults 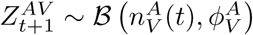 and new adults

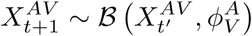

This is because right after 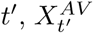 is the number of the individuals entering the adult compartment. The transitioning adults are obtained with the equation 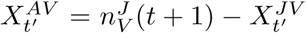as they are picked from the pool of previous juveniles. This eventually leads to (after a few simplifications)

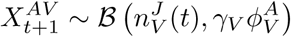

Similar arguments can be brought up for the predator. Because only the predator reproduction rate is affected by prey density at *t* in the predator submodel, and predator reproduction occurs right at *t′*, predators are assumed to depend for reproduction on prey individuals born in the previous time step.

#### Ecological consequences of the modelling assumptions

- From the predator reproduction perspective, both post- and pre-breeding models are similar, and predator depend for reproduction on individuals largely born the year before
- From the prey perspective, the post-breeding model assumes that older juvenile individuals (born the year before) suffer predation, whilst the pre-breeding model assumes that all prey, including newborns, suffer predation.
- The post-breeding model may therefore be a little more internally consistent (only the “older” juveniles affect predator reproduction and predator only affect “older” juvenile survival, not newborns). However, the pre-breeding model, where predator reproduction only depends on individuals born in the previous year, but predation also impacts newborns, is not unrealistic as well: predators that reproduce on large juvenile individuals can also eat eggs or very young chicks for their basic sustenance.

## Footnotes

1 This important step is modelled simply using a Binomial distribution because we have two stages; with more stages, it would require a Multinomial distribution, see Watkins (2000) for details

2 assuming that the year is defined by the change in year t of course, which may be equivalent to start it around January with breeding occuring early in the spring

## References

Abadi, F., Gimenez, O., Jakober, H., Stauber, W., Arlettaz, R. & Schaub, M. (2012). Estimating the strength of density dependence in the presence of observation errors using integrated population models. Ecological Modelling, 242, 1–9.

Adler, P.B. & HilleRisLambers, J. (2008). The influence of climate and species composition on the population dynamics of ten prairie forbs. Ecology, 89, 3049–3060.

Auger-Méthé, M., Field, C., Albertsen, C.M., Derocher, A.E., Lewis, M.A., Jonsen, I.D. & Flemming, J.M. (2016). State-space models’ dirty little secrets: even simple linear gaussian models can have estimation problems. Scientific reports, 6, 26677.

Barraquand, F., Louca, S., Abbott, K.C., Cobbold, C.A., Cordoleani, F., DeAngelis, D.L., Elderd, B.D., Fox, J.W., Greenwood, P., Hilker, F.M. et al. (2017). Moving forward in circles: challenges and opportunities in modelling population cycles. Ecology letters, 20, 1074–1092.

Barraquand, F. & Nielsen, O.K. (2018). Predator-prey feedback in a gyrfalcon-ptarmigan system? Ecology and Evolution, 8, 12425–12434.

Benaïm, M. & Schreiber, S. (2009). Persistence of structured populations in random environments. Theoretical Population Biology, 76, 19–34.

Besbeas, P., Freeman, S.N., Morgan, B.J. & Catchpole, E.A. (2002). Integrating mark–recapture– recovery and census data to estimate animal abundance and demographic parameters. Biometrics, 58, 540–547.

Caswell, H. (2001). Matrix populations models. Sinauer Associates Inc, Sunderland, MA.

Certain, G., Barraquand, F. & Gårdmark, A. (2018). How do MAR(1) models cope with hidden nonlinearities in ecological dynamics? Methods in Ecology and Evolution, 9, 1975–1995.

Chandler, R.B. & Clark, J.D. (2014). Spatially explicit integrated population models. Methods in Ecology and Evolution, 5, 1351–1360.

Che-Castaldo, J., Che-Castaldo, C. & Neel, M.C. (2018). Predictability of demographic rates based on phylogeny and biological similarity. Conservation Biology.

Chu, C. & Adler, P.B. (2015). Large niche differences emerge at the recruitment stage to stabilize grassland coexistence. Ecological Monographs, 85, 373–392.

Cole, D.J. & McCrea, R.S. (2016). Parameter redundancy in discrete state-space and integrated models. Biometrical Journal, 58, 1071–1090.

Cooch, E.G., Gauthier, G. & Rockwell, R.F. (2003). Apparent differences in stochastic growth rates based on timing of census: a cautionary note. Ecological Modelling, 159, 133–143.

Cushing, J. (1988). Nonlinear matrix models and population dynamics. Natural Resource Modeling, 2, 539–580.

Dennis, B., Desharnais, R.A., Cushing, J. & Costantino, R. (1995). Nonlinear demographic dynamics: mathematical models, statistical methods, and biological experiments. Ecological Monographs, 65, 261–282.

Dennis, B., Ponciano, J.M. & Taper, M.L. (2010). Replicated sampling increases efficiency in monitoring biological populations. Ecology, 91, 610–620.

Fujiwara, M., Pfeiffer, G., Boggess, M., Day, S. & Walton, J. (2011). Coexistence of competing stage-structured populations. Scientific Reports, 1, 107.

Gervasi, V., Nilsen, E.B., Sand, H., Panzacchi, M., Rauset, G.R., Pedersen, H.C., Kindberg, J., Wabakken, P., Zimmermann, B., Odden, J. et al. (2012). Predicting the potential demographic impact of predators on their prey: a comparative analysis of two carnivore–ungulate systems in scandinavia. Journal of Animal Ecology, 81, 443–454.

Gimenez, O., Morgan, B.J. & Brooks, S.P. (2009). Weak identifiability in models for mark-recapture-recovery data. In: Modeling demographic processes in marked populations. Springer, pp. 1055–1067.

Greenman, J. & Benton, T. (2004). Large amplification in stage-structured models: Arnol’d tongues revisited. Journal of mathematical biology, 48, 647–671.

Haccou, P., Jagers, P. & Vatutin, V.A. (2005). Branching processes: variation, growth, and extinction of populations. 5. Cambridge university press.

Hampton, S.E., Holmes, E.E., Scheef, L.P., Scheuerell, M.D., Katz, S.L., Pendleton, D.E. & Ward, E.J. (2013). Quantifying effects of abiotic and biotic drivers on community dynamics with multivariate autoregressive (MAR) models. Ecology, 94, 2663–2669.

Hartig, F. & Dormann, C.F. (2013). Does model-free forecasting really outperform the true model? Proceedings of the National Academy of Sciences, 110, E3975–E3975.

Hassell, M. & Comins, H. (1976). Discrete time models for two-species competition. Theoretical Population Biology, 9, 202–221.

Ives, A., Dennis, B., Cottingham, K. & Carpenter, S. (2003). Estimating community stability and ecological interactions from time-series data. Ecological Monographs, 73, 301–330.

Ives, A.R., Einarsson, Á., Jansen, V.A. & Gardarsson, A. (2008). High-amplitude fluctuations and alternative dynamical states of midges in Lake Myvatn. Nature, 452, 84–87.

Kéry, M. & Schaub, M. (2011). Bayesian population analysis using WinBUGS: a hierarchical perspective. Academic Press.

Knape, J. (2008). Estimability of density dependence in models of time series data. Ecology, 89, 2994–3000.

Kon, R., Saito, Y. & Takeuchi, Y. (2004). Permanence of single-species stage-structured models. Journal of Mathematical Biology, 48, 515–528.

Kot, M. (2001). Elements of mathematical ecology. Cambridge University Press.

Krebs, C.J., Boonstra, R. & Boutin, S. (2018). Using experimentation to understand the 10-year snowshoe hare cycle in the boreal forest of north america. Journal of Animal Ecology, 87, 87–100.

Lahoz-Monfort, J.J., Harris, M.P., Wanless, S., Freeman, S.N. & Morgan, B.J. (2017). Bringing it all together: multi-species integrated population modelling of a breeding community. Journal of Agricultural, Biological and Environmental Statistics, 22, 140–160.

Lebreton, J.D., Burnham, K.P., Clobert, J. & Anderson, D.R. (1992). Modeling survival and testing biological hypotheses using marked animals: a unified approach with case studies. Ecological monographs, 62, 67–118.

Lebreton, J.D. & Gimenez, O. (2013). Detecting and estimating density dependence in wildlife populations. The Journal of Wildlife Management, 77, 12–23.

Lebreton, J.D., Nichols, J.D., Barker, R.J., Pradel, R. & Spendelow, J.A. (2009). Modeling individual animal histories with multistate capture–recapture models. Advances in ecological research, 41, 87–173.

McKane, A.J. & Newman, T.J. (2005). Predator-prey cycles from resonant amplification of demographic stochasticity. Physical review letters, 94, 218102.

Miller, T.E. & Rudolf, V.H. (2011). Thinking inside the box: community-level consequences of stage-structured populations. Trends in Ecology & Evolution, 26, 457–466.

Murdoch, W., Kendall, B., Nisbet, R., Briggs, C., McCauley, E. & Bolser, R. (2002). Single-species models for many-species food webs. Nature, 417, 541.

Mutshinda, C.M., O’Hara, R.B. & Woiwod, I.P. (2009). What drives community dynamics? Proceedings of the Royal Society B: Biological Sciences, 276, 2923–2929.

Neubert, M.G. & Caswell, H. (2000). Density-dependent vital rates and their population dynamic consequences. Journal of Mathematical Biology, 41, 103–121.

Newman, K., Buckland, S., Morgan, B., King, R., Borchers, D., Cole, D., Besbeas, P., Gimenez, O. & Thomas, L. (2014). State-space models. In: Modelling Population Dynamics. Springer, pp. 39–50.

Nisbet, R. & Gurney, W. (1976). A simple mechanism for population cycles. Nature, 263, 319.

Péron, G. & Koons, D.N. (2012). Integrated modeling of communities: parasitism, competition, and demographic synchrony in sympatric ducks. Ecology, 93, 2456–2464.

Plummer, M. et al. (2003). Jags: A program for analysis of bayesian graphical models using gibbs sampling. In: Proceedings of the 3rd international workshop on distributed statistical computing. Vienna, Austria, vol. 124.

Preisser, E.L., Bolnick, D.I. & Benard, M.F. (2005). Scared to death? the effects of intimidation and consumption in predator–prey interactions. Ecology, 86, 501–509.

Rajala, T., Olhede, S.C. & Murrell, D.J. (2018). When do we have the power to detect biological interactions in spatial point patterns? Journal of Ecology.

de Roos, A.M. & Persson, L. (2013). Population and community ecology of ontogenetic development. Princeton University Press.

Salguero-Gómez, R., Jones, O.R., Archer, C.R., Bein, C., Buhr, H., Farack, C., Gottschalk, F., Hartmann, A., Henning, A., Hoppe, G. et al. (2016). Comadre: a global data base of animal demography. Journal of Animal Ecology, 85, 371–384.

Saunders, S.P., Cuthbert, F.J. & Zipkin, E.F. (2018). Evaluating population viability and efficacy of conservation management using integrated population models. Journal of Applied Ecology, 55, 1380–1392.

Schaub, M. & Abadi, F. (2011). Integrated population models: a novel analysis framework for deeper insights into population dynamics. Journal of Ornithology, 152, 227–237.

Travis, C., Post, W., DeAngelis, D. & Perkowski, J. (1980). Analysis of compensatory leslie matrix models for competing species. Theoretical population biology, 18, 16–30.

Tredennick, A.T., Hooten, M.B. & Adler, P.B. (2017). Do we need demographic data to forecast plant population dynamics? Methods in Ecology and Evolution, 8, 541–551.

Turchin, P. & Ellner, S. (2000). Living on the edge of chaos: population dynamics of Fennoscandian voles. Ecology, 81, 3099–3116.

Turchin, P. & Hanski, I. (1997). An empirically based model for latitudinal gradient in vole population dynamics. American Naturalist, 149, 842–874.

Valkama, J., Korpimäki, E., Arroyo, B., Beja, P., Bretagnolle, V., Bro, E., Kenward, R., Manosa, S., Redpath, S.M., Thirgood, S. et al. (2005). Birds of prey as limiting factors of gamebird populations in europe: a review. Biological Reviews, 80, 171–203.

Watkins, J.C. (2000). Consistency and fluctuation theorems for discrete time structured population models having demographic stochasticity. Journal of mathematical biology, 41, 253–271.

Weide, V., Varriale, M.C. & Hilker, F.M. (2018). Hydra effect and paradox of enrichment in discrete-time predator-prey models. Mathematical Biosciences.

Wikan, A. (2001). From chaos to chaos. an analysis of a discrete age-structured prey–predator model. Journal of Mathematical Biology, 43, 471–500.

Wood, S.N. (2010). Statistical inference for noisy nonlinear ecological dynamic systems. Nature, 466, 1102.

Zhou, C., Fujiwara, M. & Grant, W.E. (2013). Dynamics of a predator–prey interaction with seasonal reproduction and continuous predation. Ecological modelling, 268, 25–36.

Zipkin, E.F. & Saunders, S.P. (2018). Synthesizing multiple data types for biological conservation using integrated population models. Biological Conservation, 217, 240–250.

